# One Health Assessment of an Urban Temporary Settlement Reveals Gut Microbiome Serving as Antimicrobial Resistance Gene Reservoir

**DOI:** 10.1101/2023.03.15.532708

**Authors:** Rajindra Napit, Anupama Gurung, Ajit Poudel, Prajwol Manadhar, Ashok B. Chaudhary, Ajay Narayan Sharma, Samita Raut, Saman Man Pradhan, Jyotsna Joshi, Mathilde Poyet, Mathieu Groussin, Rajesh M. Rajbhandari, Dibesh B. Karmacharya

## Abstract

Antimicrobial resistance (AMR) is an emerging and growing global health challenge that could result in 10.2 million deaths annually by 2050. The unrestricted and haphazard use of antibiotics is contributing to the rapid emergence and spread of AMR, and the problem is exacerbated by release of untreated waste water from high-risk sources like hospitals into rivers. Bacteria often develop resistance through horizontal gene transfer mechanism and gut flora can act as a source for new Antimicrobial Resistance Genes (ARG). Upcoming methods like metagenomics can identify the resistance profile (AMR) of gut microbiome, and detect bacterial infections that otherwise go unnoticed. Our study focused on understanding the presence of AMR mutations and gene transfer dynamics in human, animal and environmental samples collected in one of the temporary settlements of Kathmandu (Nepal) using One Health approach. Current AMR reporting based on clinical cases is limited and does not provide information on specific pathogen and associated AMR genes-our study is an effort to contribute information to fulfill this gap.

Twenty-one samples were collected from a temporary settlement in Thapathali (Kathmandu), which included fecal samples from birds (n=3) and humans (n=14), and environmental samples (n=4). Microbiological assessment was carried out based on 16S sequence metagenomic analysis using MiSeq (Illumina, USA). Taxonomic classification on obtained 16S sequences were determined by using Metaphlan 2 and Qiime 2 bioinformatics tools. ShortBRED was used to classify ARG and virulence factors, and WAFFLE was used for horizontal gene transfer event prediction. The network analysis was carried out using Gephi v0.9 and the ResistoXplorer web tool to identify ARG in the collected samples.

*Prevotella spp*. was the dominant gut microbiome in humans. We detected diverse phages and viruses, including Stx-2 converting phages. 72 virulence factors and 53 ARG subtypes were detected, with poultry samples having the highest number of subtypes. The cluster and network analysis showed a strong association between gut microbiome and ARG, which was also supported by Horizontal Gene Transfer (HGT) analysis. One-Health interface showed ARG dynamics and revealed gut microbiomes of humans and animals serving as a reservoir for the circulating ARG.

## Introduction

Antimicrobial resistance (AMR) is an emerging global health challenge (Aarestrup, 2015). The World Health Organization (WHO) has endorsed a global action plan on AMR surveillance and strategies for mitigation (WHO, 2015). To date, drug-resistant infections are responsible for over 5 million deaths annually as per a report published in 2022 (Murray et al., 2022a) and if the looming crisis is not averted, we might see 100% drug-resistant (superbug) infections, resulting in 10.2 million deaths of working-age population by 2050 (Anderson et al., 2019; Murray et al., 2022b). Haphazard and unrestrained use of antibiotics in agricultural and health sectors have caused an emergence of new bacterial populations carrying and transferring multitude of antibiotic resistant genes (ARG) (Xu et al., 2015; Guo et al., 2017). One of the emerging sources of AMR has been non-treated or minimally treated hospital wastewater which gets released into rivers (Fouz et al., 2020), such is the case with the majority of the hospitals in Kathmandu (Nepal) (Thakali et al., 2021a). Many of the temporary settlements (slums), including our study site, have been exposed to such untreated wastewater from nearby hospitals.

Bacteria often develop AMR resistance through Horizontal Gene Transfer (HGT) and ARG are acquired from related or distant species. Mobile genetic elements (MGEs) within bacterial cells like plasmids, integrons and transposons are enhanced by recombination mechanisms like conjugation, transduction and transformation (Murray et al., 2022b). Reservoirs of ARG found in microbial communities in humans, animals and environment are crucial in propagation of AMR (Forsberg et al., 2014). Gut flora is responsible for overall health of mammalian species, including its important role in boosting host immune system and enhancing nutrition acquisition (Anthony et al., 2021). Antibiotics create a disharmony in gut ecosystems by altering their functional and taxonomic composition, enabling colonization by opportunistic pathogens (Sorbara and Pamer, 2019).

Metagenomics has become an important tool to profile AMR of gut microbiome (Hendriksen et al., 2019) and to identify various environmental niche that may be source of dissemination of AMR bacteria and resistance mechanism (Fitzpatrick and Walsh, 2016). Using next generation sequencing (NGS) data of short targeted biomarker reads, metagenomics techniques enable quantification of large number of transmissible resistance genes (Miller et al., 2013). Metagenomics is a relatively new and emerging method, and since its first published application in 2010, more cost effective NGS platforms have been developed and are now commercially available. With this technique, we can detect microorganisms without any presupposition, especially for those infections that are difficult to detect using conventional diagnostic tools (Miller et al., 2013; Schlaberg et al., 2017; d’Humières et al., 2021). This technique is particularly useful in early detection and surveillance of highly infectious zoonotic diseases (Newell et al., 2010).

Disease surveillance, including AMR, is mostly based on reports submitted by clinical or laboratory outlets (Gibbons et al., 2014). Covid-19 pandemic has highlighted the importance of a broad but accurate surveillance of communicable diseases (Assefa et al., 2022; Basseal et al., 2022). Detection and monitoring of virulence factors can be particularly useful in understanding public health risks of potential infections as virulence factors are often associated with the ability of bacteria or virus to adhere, colonize, invade and sequester nutrients from hosts (Sharma et al., 2017) and increase their pathogenicity (Miller et al., 2013). HGT is a mechanism through which bacteria evolve by transferring genetic information such as AMR, virulence and functional genes between cells (Burmeister, 2015). In this study, we have analyzed samples collected from humans, animals and environment in a high risk urban site located in Kathmandu; and detected AMR genes in bacteria, determined their virulence factors and HGT.

## Material and methods

### Study site

This study site is a temporary settlement located in Thapathali, Kathmandu (Figure). With an estimated population of 661, this settlement is situated in the banks of the Bagmati River and located in the middle of a highly dense and urbanized part of the Kathmandu valley. Two large hospitals, the Paropakar Maternity and Women’s Hospital and Norvic Hospital, are situated within 200 meters of the sampling site. Untreated wastewater from the hospitals are discharged into the nearby Bagmati river.

We collected samples from birds [n=3, fecal; chicken, *Gallus gallus domesticus*, n=1, common quails, *Coturnix coturnix*), n=2], humans (n=14, fecal), and environmental samples [n=4; water (n=2), soil (n=1), and river bed sediments (n=1)].

Ethical approval for the study was obtained from the Nepal Health Research Council (Reg. No. 792/2018).

**Figure 1:**
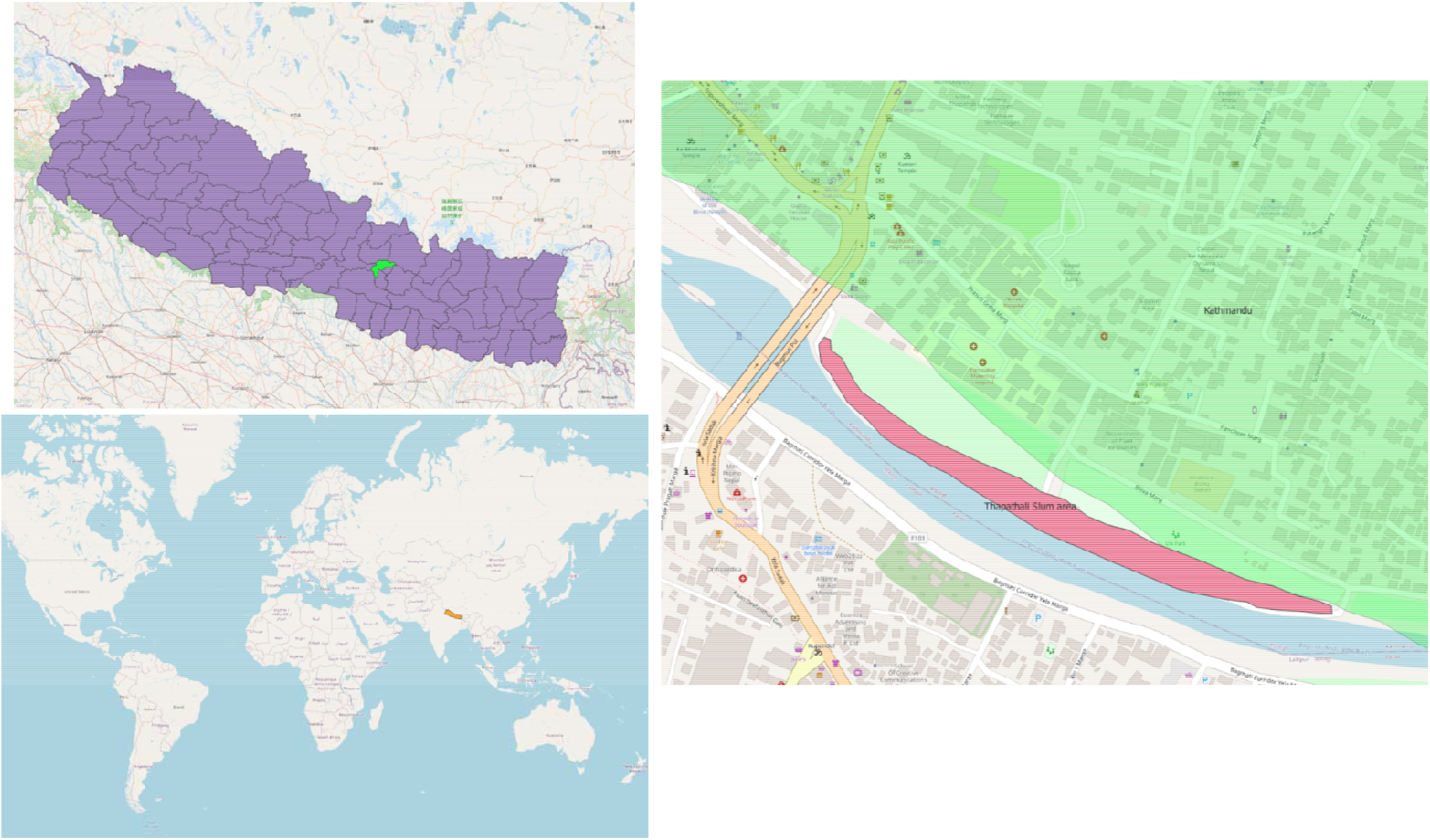
The study site-Thapathali temporary settlement, located in the Kathmandu metropolitan city. The map was created using QGIS (an open-source GIS platform) using base maps from OpenStreetMap contributors (25) and shape files from OpendataNepal.com (26)

### Sample Collection and DNA extraction

River water samples were collected in May (2019) using electric auto-sampler (Biobot Analytics Inc, USA). Five hundred milliliter (ml) gab and sediment sample were collected in zip lock bags using sterile plastic spatula. Written consent were obtained from the participating residents and structured questionnaires were used for survey. The humans and domestic animals (chicken and quails) fecal samples were collected in sterile plastic stool containers. The fecal samples were then transferred into two individual vials containing 5ml RNAlater (Thermo Fisher Scientific, USA) and Glycerol respectively and were homogenized uniformly. From 5ml of homogenized solution, 1 ml solution was transferred to each five 2ml cryovials. 1L of ground water was collected in a sterile screw capped bottle, and soil samples were collected in zip lock bags avoiding surface debris. The samples were labeled with unique identity code and GPS of site and sample collected were recorded. The samples were transported immediately to the laboratory in cold chain box maintaining temperature between 2-8° Celsius.

DNA was extracted from fecal samples using the QIAamp Fast DNA Stool Mini Kit (Qiagen, Germany) following manufacturer’s instructions. For environmental samples, DNA was extracted using the PowerSoil DNA isolation kit (MO BIO Laboratories Inc., USA). DNA concentration was measured using a Qubit™ 3 Fluorometer (Invitrogen, USA). The integrity and size of the extracted DNA were examined with electrophoresis in 0.8% agarose gel.

### 16S rRNA sequencing

16S rRNA gene was amplified using archaeal and bacterial primers (515F and 806R), targeting V3 and V4 regions (Caporaso et al., 2011). The PCR products were quantified using Qubit™ 3 Fluorometer, multiplexed at even concentration and sequenced on 600 bp (2 × 300bp) pair-end using Illumina MiSeq platform (Illumina, Inc., USA) (EMP 16S Illumina Amplicon Protocol, n.d.).

### Metagenomic library preparation and sequencing

1 ng of genomic DNA from each sample was used with Illumina MiSeq Nextera XT DNA Library Preparation Kit (Illumina, Inc., USA), the paired-end library was constructed with an insert (500 bp) for all 21 samples. DNA was cleaned by AMPure XP beads (Agentcourt, USA) and tagmented and indexed using Nextera XT Index Kit (Illumina, Inc., USA). Clean DNA was again quantified and evaluated using Qubit (Invitrogen, USA) and Agilent bioanalyzer DNA 1000 kit (Agilent Technologies, UK). Finally, all samples were pooled at 4 nm concentration and paired-end [300bp (2 × 151bp)] sequenced in MiSeq platform (Illumina, USA).

## Data analysis

### 16s rRNA bacterial taxonomic profiling

Data were analyzed using the QIIME version 2.0 pipeline. Raw sequences were de-multiplexed and then quality-filtered using DADA2 in QIIME. Sequences were then clustered into Operational Taxonomic Units (OTUs, 99% similarity) with USEARCH using the open reference clustering protocol (Amplicon analysis with QIIME2 - VL microbiome project, n.d.). The Silva_132_release database was used to assign taxonomy and resulting OTU table was then rarefied based on alpha rarefaction of 21,383 reads per sample.

### Metagenomic taxonomic profiling

The metagenomic phylogenetic analysis was done using tool-MetaPhlAn V 3.0 (https://github.com/biobakery/MetaPhlAn). The analysis was performed as described in the manual of MetaPhlAn V 3.0 to run an analysis (https://github.com/biobakery/MetaPhlAn/wiki/MetaPhlAn-3.0) (MetaPhlAn 3.0 ·biobakery/MetaPhlAn Wiki ·GitHub, n.d.).

### Virulence factor and antimicrobial resistance gene (ARG) analysis

The shotgun sequence data was used to analyze AMR and Virulence factor (VF) gene using tool ShortBRED (Kaminski et al., 2015). Antibiotic Resistance Database (ARDB) markers and Virulence Factors Database (VFDB) markers were used in the analysis, which is the default database present in ShortBRED tools. Similar to the MetaPhlAn, manual of ShortBRED was used to perform the analysis (https://github.com/biobakery/shortbred) (GitHub - biobakery/shortbred: ShortBRED is a pipeline to take a set of protein sequences, reduce them to a set of unique identifying strings (“markers”), and then search for these markers in metagenomic data and determine the presence and abundance of the, n.d.). Antimicrobial abundance data were visualized using ResistoXplorer web platform (Dhariwal et al., 2021).

### Horizontal gene transfer (HGT) Analysis

The horizontal gene transfer event was predicted using the tool WAAFLE [43] in default taxonomy database and following the prescribed manual (https://github.com/biobakery/waafle) (GitHub - biobakery/waafle: WAAFLE (a Workflow to Annotate Assemblies and Find LGT Events) is a method for finding novel LGT (Lateral Gene Transfer) events in assembled metagenomes, including those from human microbiomes., n.d.).

### Network analysis

Network analysis was performed on AMR relative abundance data obtained through ShortBRED and loaded to Gephi V 0.92-association between detected ARG in various assessed samples were then visualized. Additionally, Association Network analysis was obtained using integration analysis of AMR and taxonomic data on the ResistoXplorer Web platform. The analysis was carried out under sequence abundance cutoff=10%, correlation co-efficient cutoff=0.6, adjusted P=0.05 and 1000 permutations.

**Figure 1:**
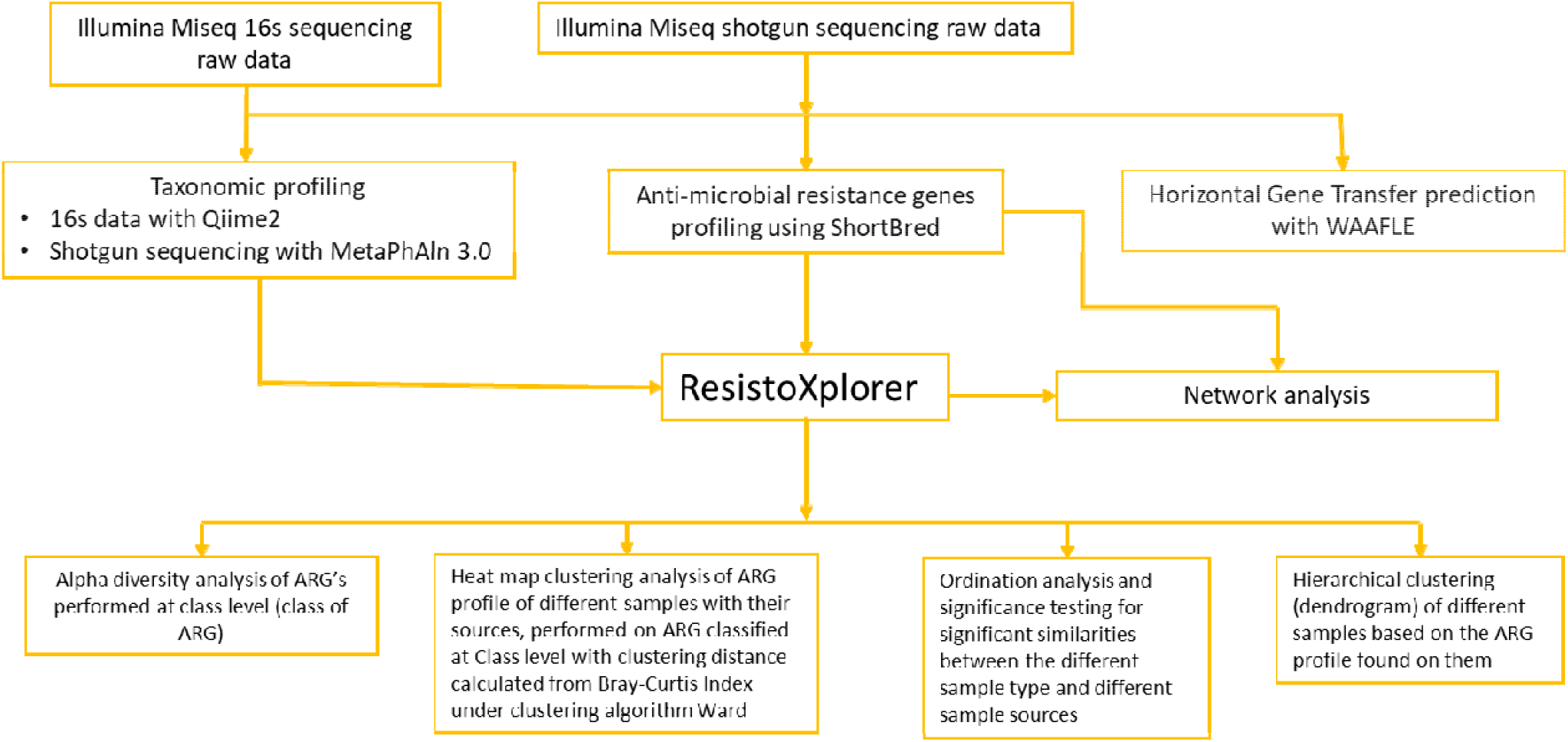
Bioinformatics data analysis workflow used in this study to determine bacterial taxonomic profiles, virulence factors, Antimicrobial Resistance Genes (ARG), Horizontal Gene Transfer (HGT) and AMR associated network analysis.

## Results

Only 11 samples (8 humans and 3 birds fecal samples) yielded 16s rRNA sequencing data. Our Shotgun metagenomics approach generated 29,000 to 2.1 million reads per sample. However, one sample only had few hundred reads (water sample-EW70) and was considered a sequencing failure. All raw data was submitted to the NCBI under Bio-project PRJNA881338.

### 16s rRNA bacterial and metagenomic taxonomic profiling

We identified various genera of bacteria (Supplementary table 1) and phages (Table **1**). Taxonomic classification of bacterial phylum rank showed dominance of phylum *Firmicutes* and *Bacteroidetes* in human samples, *Firmicutes* and *Proteobacteria* in fowl samples and *Bacteroidetes* and *Proteobacteria* in the environmental samples (Supplementary Table 2).

**Table 1:** Taxonomic profile of viruses (phage) and their relative abundance of ≥10 in samples obtained through metagenomic sequencing.

| Phage name | Family | Order | Sample | Occurrence in sample (no.) |
| --- | --- | --- | --- | --- |
| Acholeplasma_virus_L2 | Plasmpviridae | Caudovirales | Environment | 1 |
| Bacteroides_phage_B124_14 | Siphoviridae | Caudovirales | Human | 1 |
| Bacteroides_phage_B40_8 | Siphoviridae | Caudovirales | Human | 1 |
| Enterobacteria_phage_cdtI | Siphoviridae | Caudovirales | Human | 1 |
| Enterobacteria_phage_P4 | Caudovirales unclassified | Caudovirales | Human | 1 |
| Enterobacteria_phage_YYZ_2008 | Siphoviridae | Caudovirales | Human | 1 |
| Escherichia_phage_KBNP21 | Podoviridae | Caudovirales | Human | 1 |
| Escherichia_phage_TL_2011b | Podoviridae | Caudovirales | Human | 1 |
| Escherichia_virus_K30 | Podoviridae | Caudovirales | Human | 1 |
| Escherichia_virus_P1 | Myoviridae | Caudovirales | Human & Poultry | 2 |
| Escherichia_virus_phiV10 | Podoviridae | Caudovirales | Human | 1 |
| Escherichia_virus_wV8 | Myoviridae | Caudovirales | Poultry | 1 |
| Klebsiella_phage_JD001 | Myoviridae | Caudovirales | Human | 1 |
| Klebsiella_virus_KP32 | Podoviridae | Caudovirales | Human | 1 |
| Lactococcus_phage_P087 | Siphoviridae | Caudovirales | Human | 1 |
| Planktothrix_phage_PaV_LD | Podoviridae | Caudovirales | Environment | 1 |
| Salmonella_phage_Vi_II_E1 | Siphoviridae | Caudovirales | Environment | 1 |
| Salmonella_virus_Epsilon15 | Podoviridae | Caudovirales | Human | 1 |
| Staphylococcus_phage_StB20 | Siphoviridae | Caudovirales | Poultry | 1 |
| Streptococcus_virus_7201 | Siphoviridae | Caudovirales | Human & Poultry | 2 |
| Streptococcus_virus_DT1 | Siphoviridae | Caudovirales | Human | 1 |
| Streptococcus_virus_phiAbc2 | Siphoviridae | Caudovirales | Human | 1 |
| Streptococcus_virus_Sfi21 | Siphoviridae | Caudovirales | Human | 1 |
| Stx2_converting_phage_1717 | Siphoviridae | Caudovirales | Human & Poultry | 5 |

### Bacteria profile in various samples

Genus *Prevotella and Escherichia* were the most prevalent bacteria found in the human samples. Other bacterial species present belonged to genus *Lachnospira, Roseburia, Eubacterium, Faecalibacterium, Bacteroides* and *Butyrivibrio* (Supplementary Table 1). 16s data revealed genus *Agathobacter, Bacteroides, Prevotella, Escherichia, Clostridium, Streptococcus, Blautia, Lachnospira, Feacalibacterium, Dorea* and *Roseburia* were abundant, along with presence of more than 50 other bacteria genera in the human samples (Supplementary Table 2). According to both 16s and shotgun sequencing data, bacterial population showed more variance in human samples than birds and environmental samples. Bird samples were dominated by genus *Lawsonia, Escherichia, Gallibacterium, Helicobacter and Chlamydia*, in addition to the presence of other genera (Supplementary: Table 1 and 2). Environmental samples were dominated by presence of some well characterized environmental bacterial genus such as *Pseudomonas, Aeromonas, Acinetobacter*, and *Acrobacter* (Figure 2).

**Figure 2:**
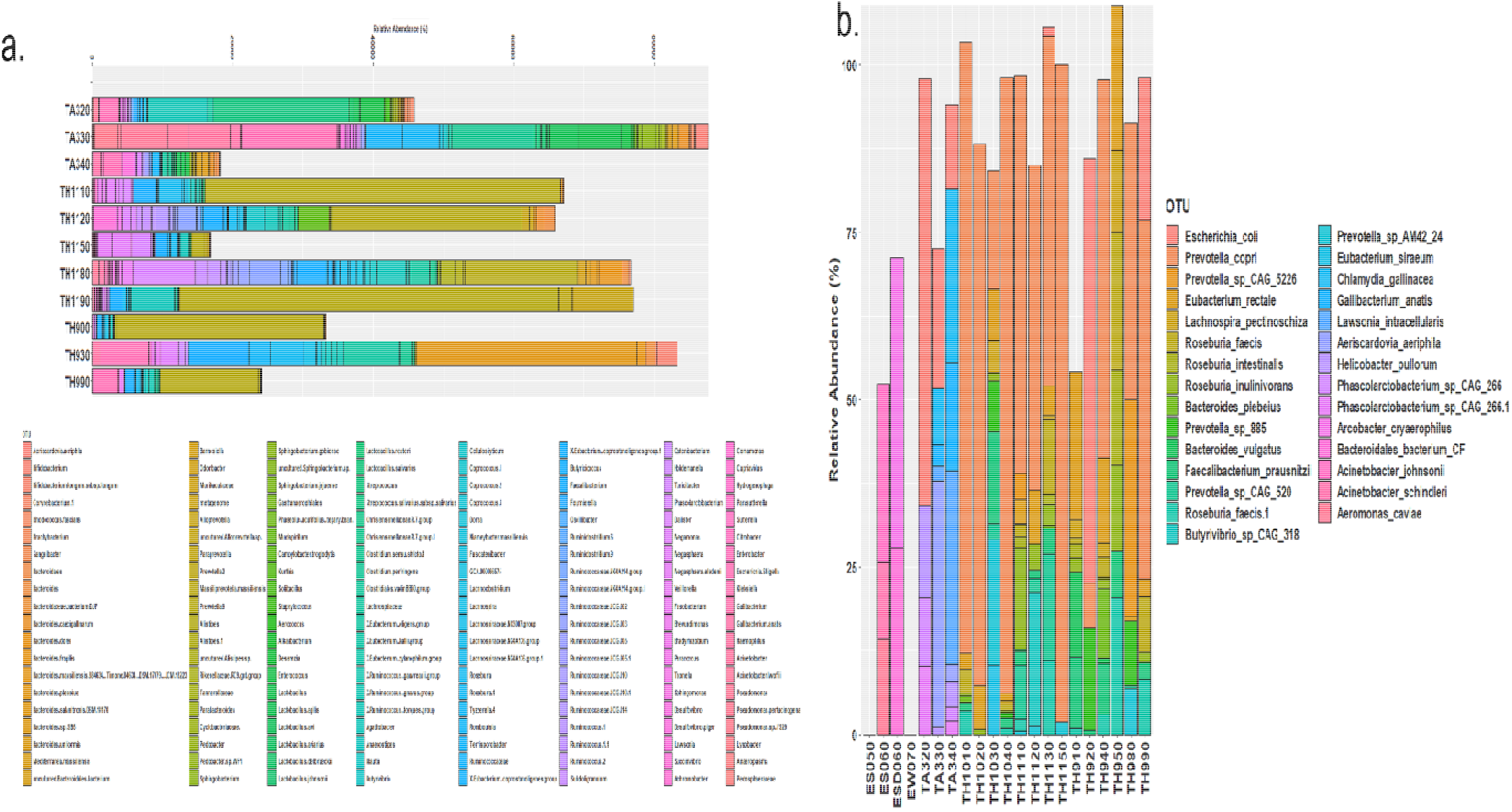
Detected bacterial phylum in various samples based on-A) 16s rRNA sequencing and B) Shotgun sequencing obtained from MetaPhlan V2.0. The plots were generated using ggplot2 in R studio 2022.07.1 Build 554

Besides these gut microbiomes, bacteria associated with human health such as *E. coli, Campylobacter, Shigella* and *Haemophilus* were also detected. Poultry pathogens-*Chlamydia gallinacea, Gallibacterium anatis* and *Helicobacter pullorum* were found in poultry samples. Aside from pathogenic organisms, known probiotic organisms such as *Lactobacillus johnsonii, Lactobacillus agilis, Lactobacillus reuteri* and *Lactobacillus salivarius* were also found in the bird samples.

### Taxonomic profile of Virus (Phages)

Taxonomic classification of virus family revealed the dominance of family *Siphoviridae*, followed by *Podoviridae* (Figure 3), both belonging to the order *Caudovirales*. Majority of the viruses were various phages of *Enterobacteriaceae* family-*Escherichia phage, Salmonella phage, and Klebsiella phage* (Table **1**). Phage diversity were highest in the human samples compared to poultry and environmental samples. In birds and humans, *Escherichia* phage along with Stx-2 converting phage that carries shiga toxin genes were found in greater numbers compared to the environmental samples. In environmental samples, *Planktothrix* phage PaV LD, *Lactococcus* phage P087 and *Achloplasma* virus L2 that infect environmental bacteria were abundant. Furthermore, in human samples, as expected phages of gut microbiome and gut pathogen were abundantly detected (Table **1**).

**Figure 3:**
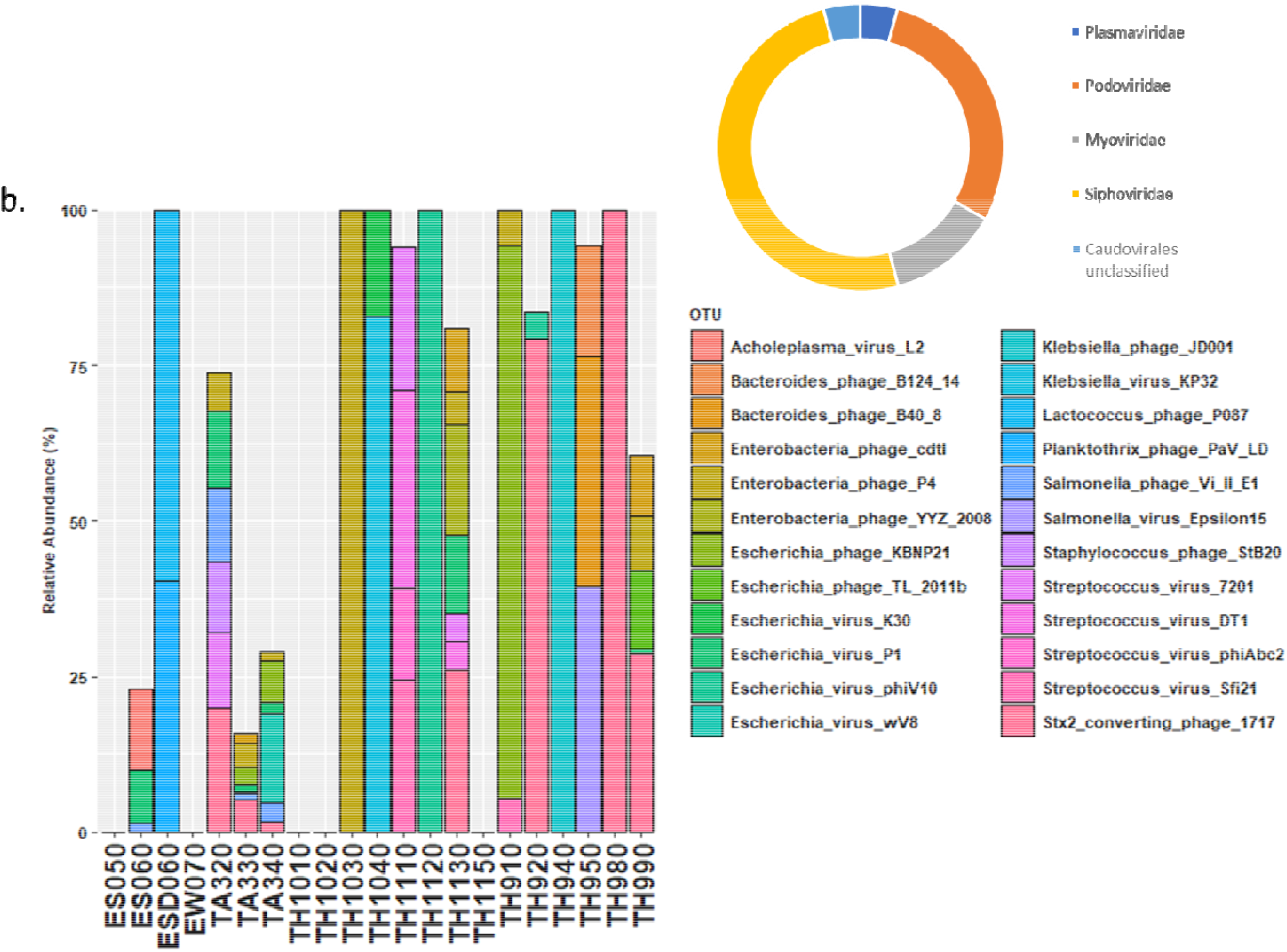
Prevalence of virus family in the samples obtained from metagenomic sequencing data. A) Family wise distribution of viruses (phage) detected B) Various phages detected in different samples; The Bar plot was generated using ggplot2 in R studio 2022.07.1 Build 554 with R version 4.2.0.

### Virulence factors profiling

Overall, a total of 72 genes that code for virulence factor were detected by shotgun metagenomics (threshold of 99 % identity and relative abundance of ≥10). Genes coding for toxins, type I to VI secretion systems, regulatory proteins, adhering proteins, siderophores as well as polysaccharides that compose the capsules and exhibit anti-phagocytic properties were detected. Virulence factors were detected predominantly from human samples, followed by poultry samples. *pilT* gene of *Pseudomonas aeruginosa* was the only gene recovered from the environmental sample (Supplementary 4). Most of the virulence genes belonged to *Escherichia coli* followed by *Shigella dysenteriae, Yersinia pestis* and *Salmonella enterica*. Very unique and specific genes were detected in some bacterial species-*Shigella flexneri, Pseudomonas aeruginosa, Legionella pneumophila* and *Yersinia enterocolitica*.

We detected genes associated with the toxigenic *E. coli* and *Shigella flexneri* exclusively in the human samples. These genes include *sat1, ltb, lta, astA* and *senB* which code for aecreted auto transporter toxin of *Enterobacteriaceae*, enterotoxin of *Enterotoxigenic Escherichia coli* (ETEC), heat stable enterotoxin of *Enteroaggregative E. coli*, and enterotoxin 2 of *Shigella flexneri* respectively (Table *2*).

**Table 2:**
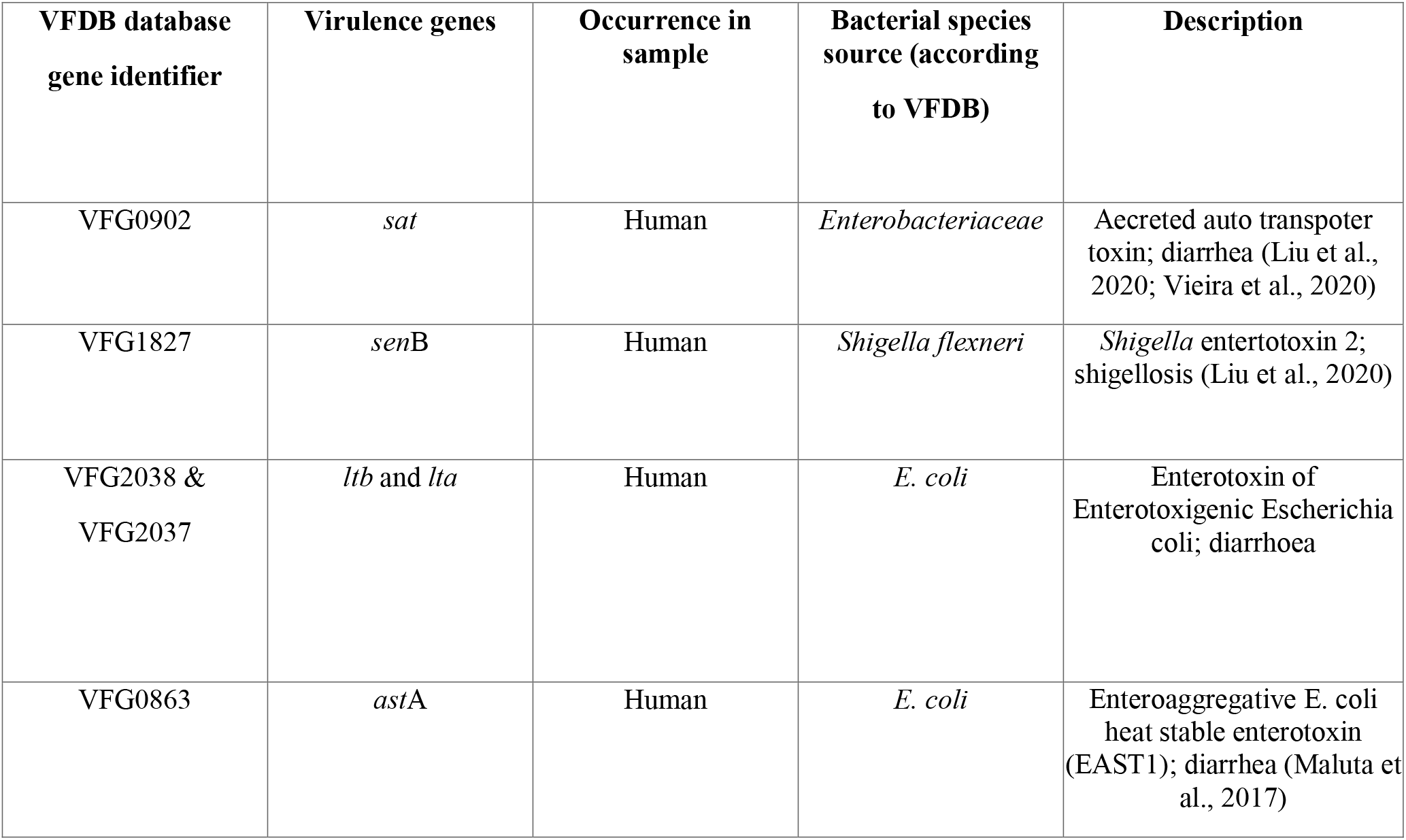
Detected toxin coding genes associated with bacteria as obtained through ShortBRED tool.

### Antibiotic resistance determinants

A total of 25 various classes of AMG and 53 ARG subtypes were detected in the 21 samples. Among these, two of the genes-*tetQ* and *ermF* were present in 14 out of 21 samples-mainly in the human and bird samples. Except *tetQ* and *ermF, tet(W)* (13/21), *cfxA* (11/21), *tet(40)* (10/21), *tet(0)*(10/21) and *tet32* (8/21) all other ARG were abundantly present. The bird samples (n=37) had the highest number of ARG, followed by the human samples (n=27) and the environmental samples (n=16).

Human, bird and environmental samples all had mobile genetic element-*intl1*. Genes responsible for resistance to antimicrobial agent such as fluoroquinolone (*qnrB6, qnrS1, QRDR*), sulfonamide (*sul1, sul2, sul3*), macrolide (*mphK, macB, macA_3, ereA2, ermCd, ermQ, ermG, ermF, ermGT, ermB*), lincosamide (*inuB*), kausagamycin (*ksgA*), vancomycin (*vanR*), undecaprenyl pyrophosphate (*bacA*), trimethoprim (*dfrA, dfrXV, dfrA14*), chloramphenicol (*catA, catQ, floR*), polymyxin (*arnA*), aminoglycosides [*aacC3, aac(60)-Ie-aph(200)-Ia, aadA, aph6, ant(3”), ant (4), ant (6), aadB, aadE, aph(3’)-IIIa, aph(3’)-Ib*], tetracycline (*tetC, tet39*, t*etQ, tet32, tetW, tetM, tet40, tetA, tetL, tetB)* and betalactam (*cepA, cfxA, bla*_TEM_, *bla*_VEB-1,_ *bla*_CTX-M_, *bla*_EC_) antibiotics were detected.

Among all the ARG subtypes, genes resistant to aminoglycosides, tetracycline, and beta-lactams were predominant. In human samples, the dominant ARG belonged to tetracycline resistance genes (*tetQ, tet32, tetW, tetM, tetO*), followed by beta-lactam resistance genes (*cepA, cfxA, bla*_CTX-M_, *bla*_EC_) and aminoglycoside resistance genes (*aadE, aph(3’)-Ib, aph6*). In bird samples, the dominant ARG were aminoglycoside resistance genes [*aacC3, aac(60)-Ie-aph(200)-Ia, aadA, ant (3”), ant (6), aadB, aadE, aph(3’)-IIIa, aph(3’)-Ib*], followed by tetracycline resistance genes (*tetQ, tet32, tetW, tetM, tet40, tetA, tetL, tetB*), and erythromycin resistance genes (*ermB, ermF, ermG*). The environmental samples had high presence of sulphonamides (*sul1, sul2, sul3*) and tetracycline [*tetM, tetA(39), tetC*]. Despite having the least number of samples, bird (poultry) samples had the highest number of ARG, while the environmental samples had the least number of ARG present. (Table *3* and Figure **4**).

**Table 3:**
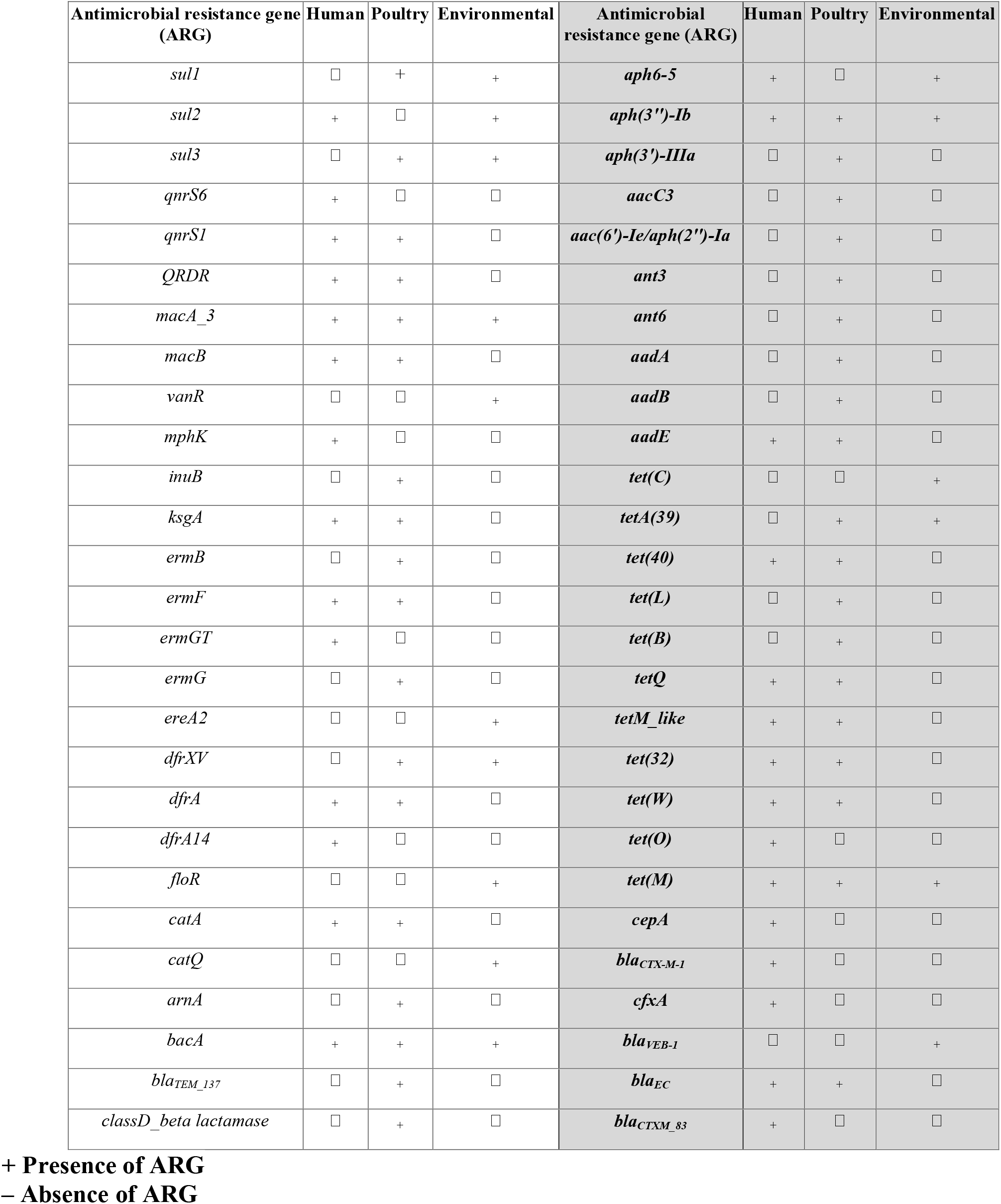
Antimicrobial resistance Genes found in human, bird (poultry) and environmental samples.

**Figure 4:**
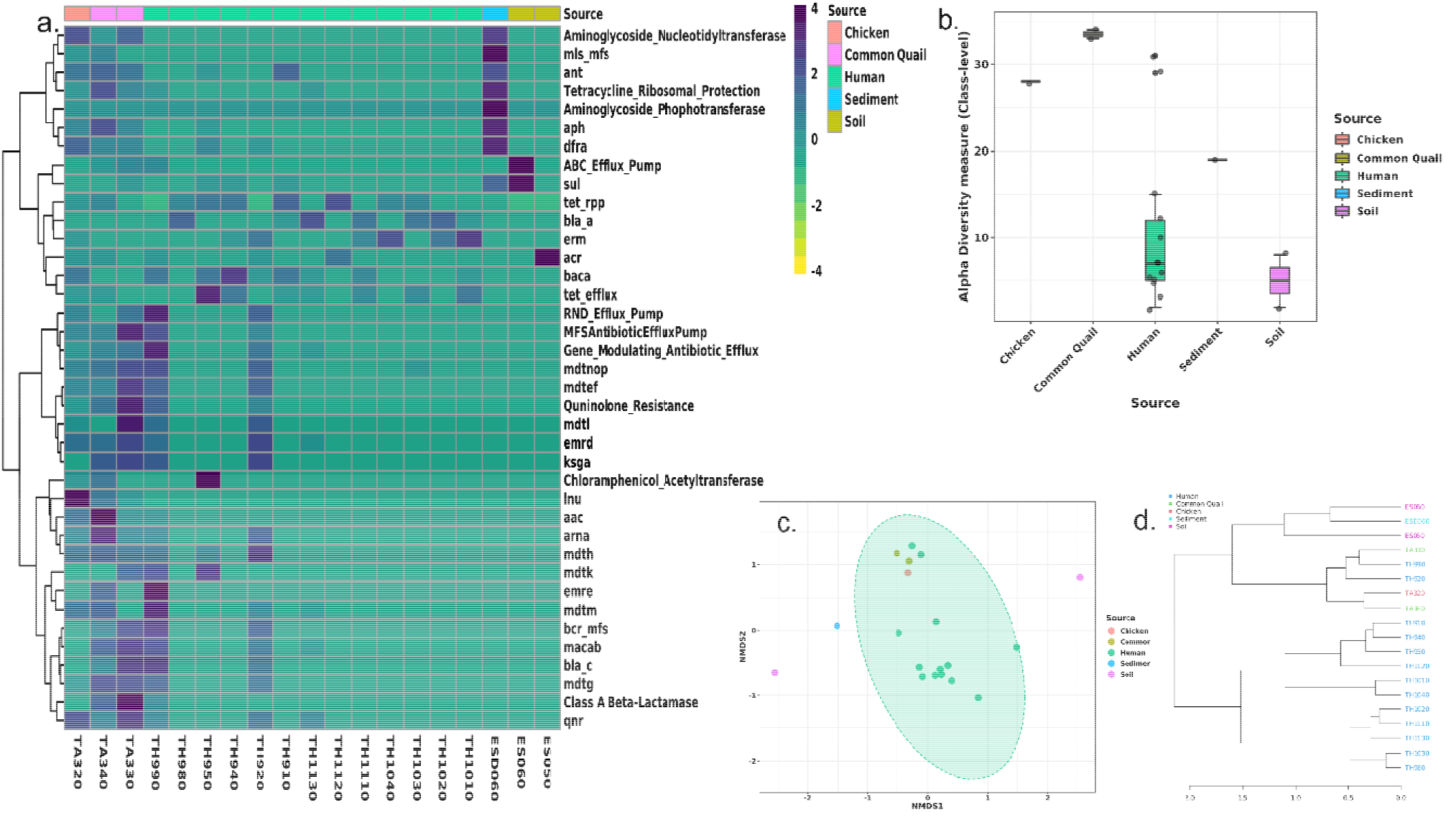
Visualization of ARG profile of various samples obtained from shot-gun sequencing processed in SHORTBred pipeline. a) Heat map obtained through cluster analysis of ARG profile of various samples and their sources, performed in ARG classified at Class level with clustering distance calculated from Bray-Curtis Index under clustering algorithm Ward. b) Alpha diversity analysis of ARG at class level, measuring diversity with Shannon. The ARG diversity is significant (P-value= 0.019166) tested using ANOVA. c) Ordination analysis and significance testing for significant similarities between the different sample types and sources. Non-metric multidimensional scaling (NMDS) ordination method was used with Bray-Curtis Index and statistically tested with Permutation MANOVA (PERMANOVA) (F-value=2.187, R-squared=0.38456 and P-value=<0.003). Samples with statistically significant ARG association are highlighted with light green shade. d) Hierarchical clustering (dendrogram) of various samples based on the ARG profile. The clustering was performed with algorithm Ward and Bray-Curtis Index as a distance measure algorithm.

The most abundant ARG in some samples (TA340, TA330, TH990, and TH920) are visualized in the generated heat-map (Figure **4**a). The highest diversity was observed in TA340 and TA330. As seen in hierarchical clustering (dendrogram) (Figure **4**d), the ARG were confined to four main clusters: two clusters were entirely from human samples, one from environmental samples and one had ARG common in the human and bird (poultry) samples. ARG found in human samples had positive association/correlation except for TH990 and TH920.

Association of ARG in birds and humans were significant as seen in the fourth cluster (F-value=2.187, R-squared=0.38456 and P-value=<0.003) (Figure 5c). ARG from the environmental samples, on the other hand, appear to be clustering within themselves without any correlation with other samples. The most diverse ARG were seen in common quail samples, while soil samples had the least number of ARG (Figure 5b). The highest variability of ARG were detected in the human samples (Figure **4**b).

**Figure 5:**
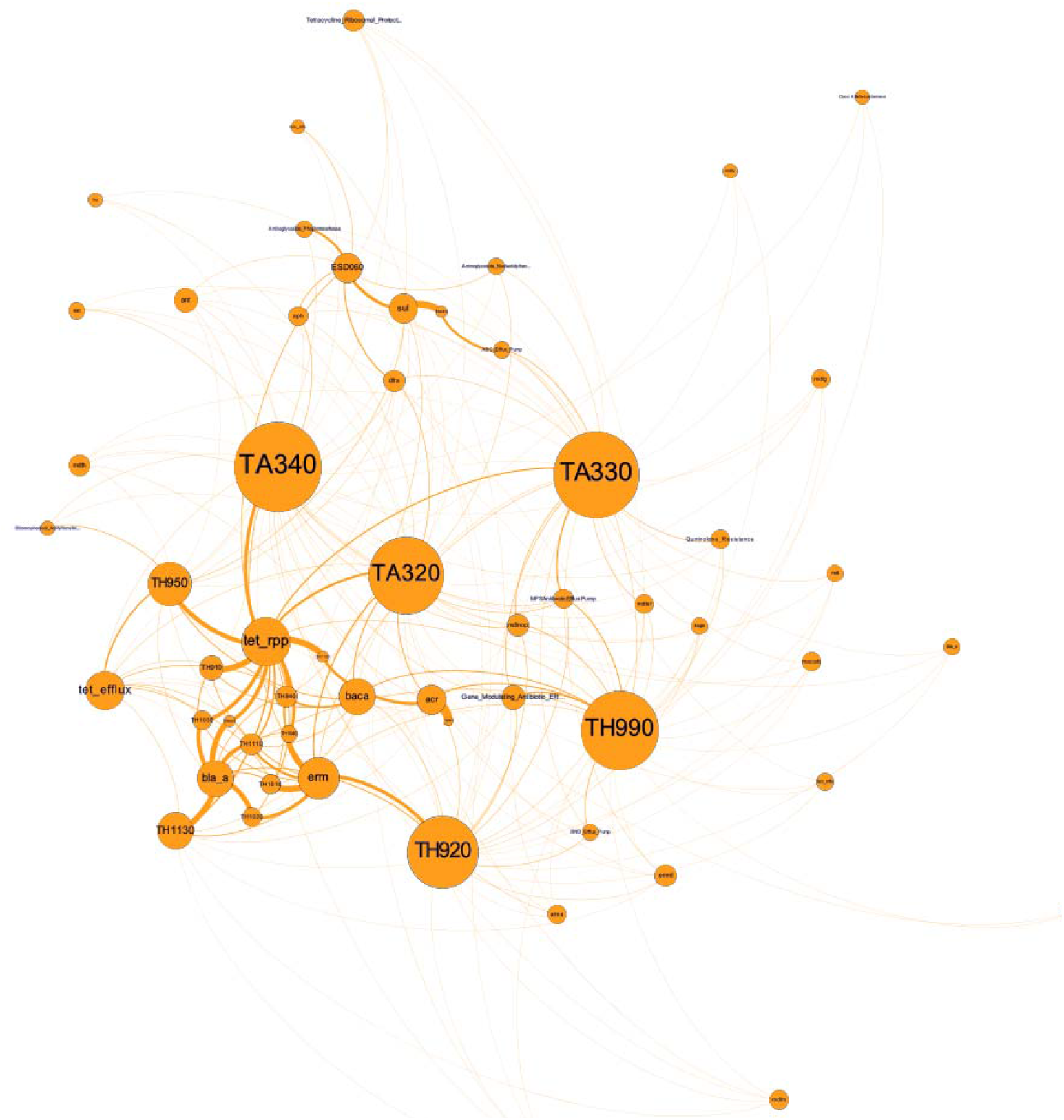
Association network analysis of ARG in various samples. The analysis was carried out with abundance data with associated metadata file on Gephi V0.92. The size of the circle represents number and strength of association-larger circle means a greater number of ARGs in the sample and also has more association with ARGs and other samples. The thickness of connections indicates strength of the association.

### Horizontal gene transfer (HGT) events

Environmental bacteria exhibited transference of genes that code for translational enzymes and RNA-directed DNA polymerase, including genes such as copper resistance proteins B that help in survival on harsh environmental conditions. Furthermore, genes that code for enzymes such as integrase and transposase of IS elements of family (*IS3/IS911* and *IS1595)* that might help them in lysogenic transformation (Antimicrobial resistance, n.d.; Davies and Davies, 2010) were also found along with genes that code for antimicrobial resistance, transcription regulation, DNA methyltransferase, and ATPases (Hamner et al., 2019).

HGT were also detected in the bird samples-especially with genes that code for proteins of replication and translation, as well as genes associated with enzymes of various pathways (mannonate dehydratase, oxidoreductase, peptidyl-prolyl cis/trans isomerases and NADH-azoreductase). In bird (poultry) samples, integrases and transposons of the *IS66* and *IS21* families that help in AMR transference (Hjelmsø et al., 2019) were detected. Clindamycin resistance transfer factor BtgB (Supplementary table 3 and Table 5), which is required for conjugal transfer of the clindamycin resistance gene in Bacteroides species (Das et al., 2020) were also detected.

In human samples, most of the commensal bacterial species HGT were detected that code for proteins of replication, transcription, translation, ion transportation, enzyme regulation, enzymes for various constitutive and inducive pathways, and enzymes required for bacterial quotidian functions (Supplementary Table 3). Various genes that code for AMR mechanisms such as ABC multidrug transport system, multidrug resistance protein (MATE family), VanY domain containing protein, penicillin binding protein (PBP) 1A, aminoglycoside phosphotransferase, *tetR* protein, and metallobetalactamase domain protein (Table 4) were detected circulating in gut microbiome. Additionally, ARG that facilitate movement and insertion of these genes were also detected. Genes associated with integrase and transposase of *IS116/IS110/IS902, IS30, IS605, IS200*, and *IS4* families were found in the gut commensals (Supplementary table 3 and Table).

**Table 4:**
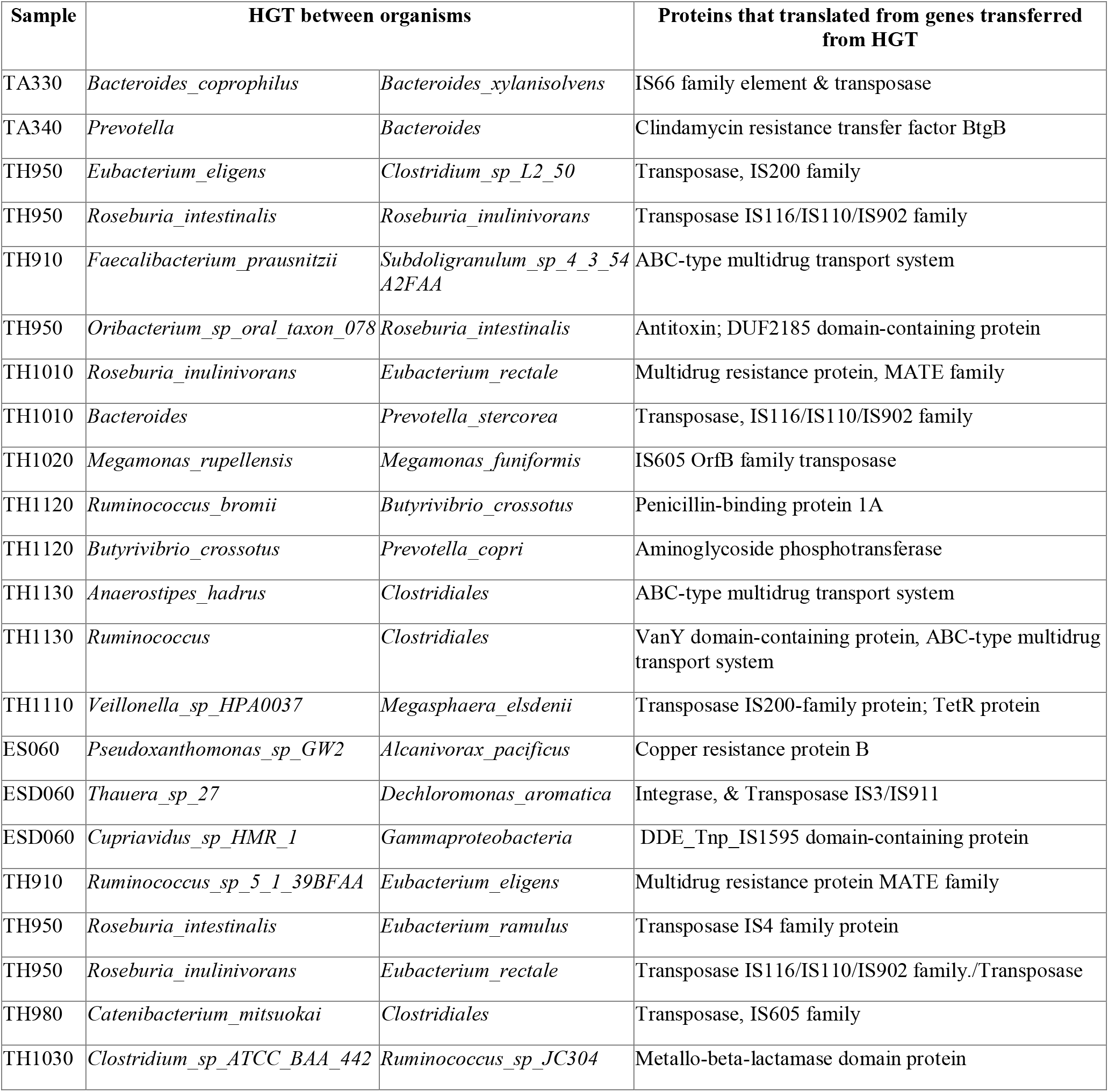
Various antimicrobial resistance proteins, virulence protein, transposase and integrase that translated from genes transferred via HGT between organisms.

### ARG Network Analysis

#### Association of ARG types in samples based on relative abundance of genes

The association network analysis of ARG in samples (Figure 6) showed strong relationship between the environmental samples (ES050, ES060, and ESD060) and aminoglycoside phosphotransferase, aminoglycoside nucleotidyltransferase, sul gene, dfrA gene, acr gene, and ABC efflux pump. In contrast, bird (poultry) samples (TA340, TA330, and TA320) showed a strong association with the tetracycline resistance gene (*tet*). Tetracycline resistance gene and tetracycline efflux had the highest association with most samples. Human samples had strong associations with *bacA, erm, acr, tet, tet* efflux, and class A betalactamase. Samples TA340, TA330, TA320, TH950, and TH920 had the highest number of ARG genes, as seen in Figure 6.

**Figure 6:**
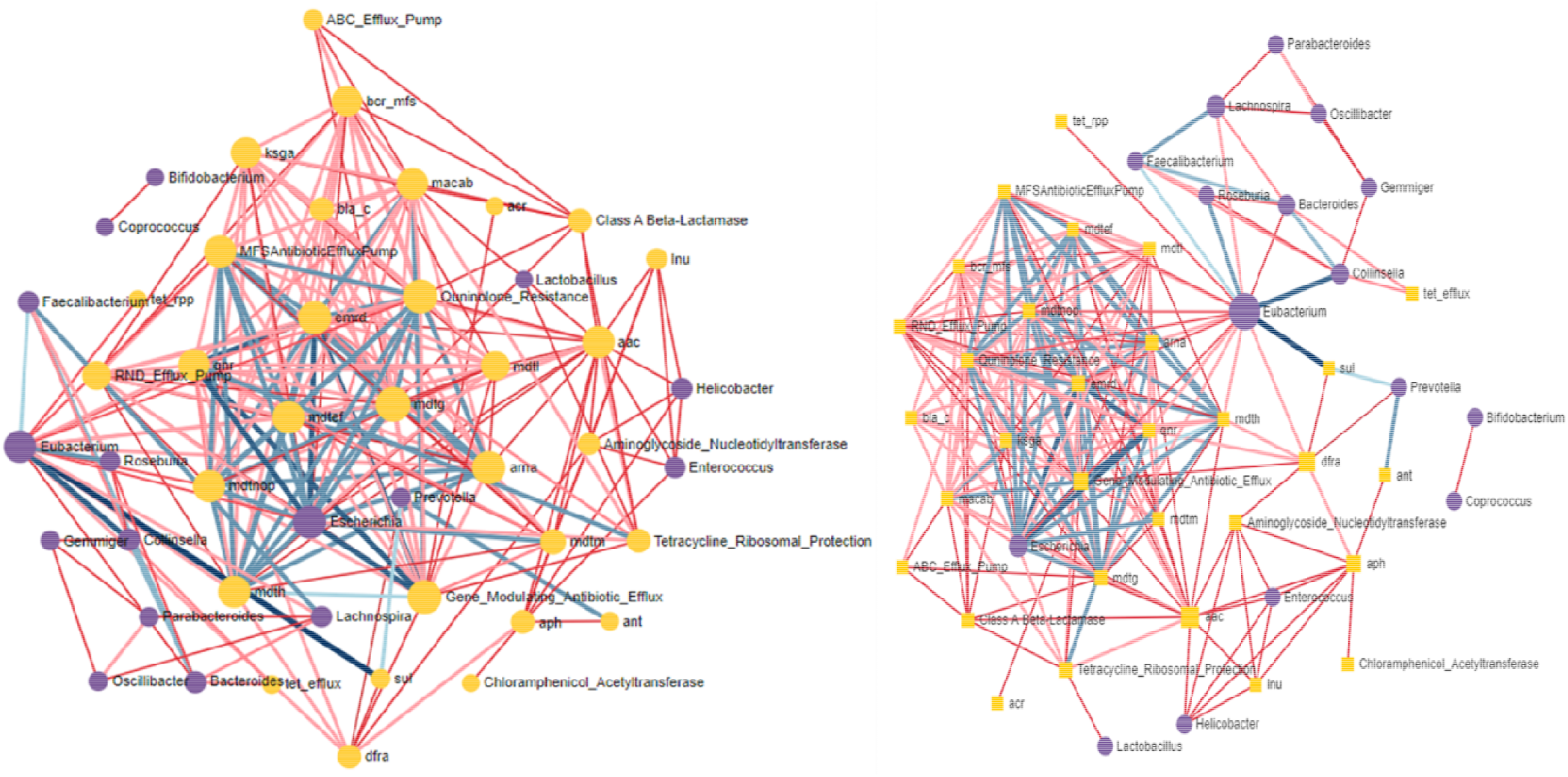
Integrative MIC analysis of ARG and Taxonomic profile of various samples using ResistoXplorer web platform. The color and shape of nodes are based on the type of data or profile (Resistome: yellow square; Microbiome: purple circle). In the network, the size of node represents the network centrality-based measure (degree or betweenness). While the color and width of edges shows the strength of correlation between them (MIC: varies from 0 to 1). A) Node size is based on Degree while B) Node size based on betweenness.

#### AMR and taxonomic profile integrative association network analysis

Connection between all detected bacteria (at genus level) and ARG (at class level) were assessed using the MIC (Maximal Information Coefficient) correlation coefficient. In network analysis (Figure 6), only the strong connections between bacteria to bacteria, bacteria to ARG, and ARG to ARG were visualized. The strong connections represented MIC values corresponding to pre-calculate *p*-values (with *p* < 0.05) based on the total number of samples. Network analysis showed the strongest associating bacteria a *Eubacterium* and demonstrated strong association between *Eubacterium* and *Faecalibacterium, Eubacterium* and *Roseburia, Eubacterium* and *Collinesella, Faecalibacterium* and *Lachnospira, Feacalibacterium* and *Bacteroides*. In ARG, the strongly associating ARG seem to be *qnr, erm*D, *arn*A, and *aac*, among the different genera of bacteria. Furthermore, efflux pumps were also found to be present in a high number and seemed to be strongly associated with *Escherichia. Escherichia* seems to be a hub for most ARG and efflux pumps and almost all AGR are strongly related to genus *Escherichia (*Figure 6).

## Discussion

AMR has become one of the most pressing issues of public health concern of the 21^st^ century (Prestinaci et al., 2015). Rise of new pathogen (bacteria or viruses) that may lead to emerging and re-emerging diseases, needs to be surveilled and understood properly (Kelly et al., 2020). Continuous misuse of antibiotics can lead to multidrug-resistant bacteria or superbugs (Antimicrobial resistance, n.d.). AMR often develop with same molecular mechanism as emergence of new bacteria (Davies and Davies, 2010; Bartoli et al., 2016). Recently, utility of metagenomics tool in epidemiological and environmental studies have been growing (Hamner et al., 2019; Hjelmsø et al., 2019; Das et al., 2020). We have used metagenomics approach to investigate AMR presence and transfer dynamics in environmental samples from One Health perspective.

Taxonomic profile obtained through shotgun sequencing revealed the genus *Prevotella* as most prevalent bacteria, especially in the human samples (Supplementary Table 1 and Table 2). According to Wu et al. 2011, *Prevotella* spp. are SCFA (short chain fatty acids) producers and can be seen in healthy individuals consuming high carbohydrate and simple sugars diets (Wu et al., 2011). Since, Nepalese diet is mainly carbohydrate based, this might explain the dominance of *Prevotella* spp.

Pathogenic bacteria such as *Shigella, Campylobacter, Haemophilus* and *E. coli* were detected in various samples. Virulence genes facilitate the organism to colonize, survive and cause diseases (Thomas and Wigneshweraraj, 2014). Even a single gene can enhance the pathogenicity- for example, presence of Shiga toxin coding gene in *E. coli* causes bloody diarrhea and hemolytic uremic syndrome (HUS) (Sandhu and Gyles, 2002). Detection of virulence genes can often indicate the presence of certain pathogens- such as *sen*B gene *for Shigella flexneri, csgG* and *rpoS* genes for *Salmonella enterica* var typhimurium and *fliR* gene for *Yersinia enterocolitica* (Supplementary table 4). Furthermore, detection of various virulence genes of *E. coli* such as *astA, ltb*, and *lta* (Table 3) can explain the presence of *enterotoxigenic E. coli* (ETEC) and *enteroaagregative E. coli* (EAEC). Detecting these genes through metagenomics can indicate the presence of various pathogens which are often missed to be detected using conventional diagnostic tools (Miller et al., 2019).

*Chlamydia gallinacea* which causes slow growth and reduced body mass in chickens was detected in two poultry samples collected from the backyard farms. Infection with this organism is also known to cause mild symptoms in humans as well (Salas and de Vega, 2008; Tariq et al., 2020). Additionally, *Helicobacter pullorum* which causes enteritis in poultry and zoonotic colitis in humans was also detected, along with *Gallibacterium anatis* that impacts egg production by triggering oophoritis, salpingitis and peritonitis in hens (Narasinakuppe Krishnegowda et al., 2020; Crisci et al., 2021). These organisms have yet to be reported in poultry in Nepal. These findings indicate many yet to be detected and identified pathogens that are floating around in our environment.

The viral families detected in this study were mostly *Siphoviridae, Podoviridae* and *Myoviridae*. Since *enterobacteria* such as *E. coli, Klebsiella, Shigella*, and gut bacteria like *Bacteroides, Prevotella, Roseburia, Lachnospira* were mostly present in the samples, phages that infect those bacteria were also found. Most of the phages that infects genera of *Enterobacteria, Prevotella and Bacteroides* fall into the families of *Siphoviridae* (Tariq et al., 2020; Thakali et al., 2021a), *Podoviridae* (Manandhar et al., 2020) and *Myoviridae* (Salas and de Vega, 2008; Crisci et al., 2021). The Stx-2 converting bacteriophage (Table 2) is phage of much importance. It induces Shiga toxin producing *E. coli* (STEC), which causes diarrhea and hemolytic uremic syndrome (HUS) in children. STEC produces Shiga toxin and genes encoding this Shiga toxin are located in the genomes of Stx-2 converting phages which are transferred by the lysogenic cycle (Acharya et al., 2017; Joshi et al., 2017). This phage was identified in seven samples (human and poultry). *E. coli* was detected in every sample, which implicates the probability of STEC production in many of these samples (Miller et al., 2019). Interestingly, phages and bacterial diversity were very similar in most of the samples, with human samples containing highest diversity (Table 1 and Supplementary table 1 and 2). This indicates that presence of any bacteria in an ecosystem, most likely harbors associated phages as well (Nayaju et al., 2021).

In our study numerous ARG subtypes (n=53) were detected. Some of these genes have been previously detected in Nepal while others like *inuB, catQ, ksgA, floR*, and *bla*_EC_ (Table 3) were detected for the first time (Acharya et al., 2017; Joshi et al., 2017; Manandhar et al., 2020; Thakali et al., 2021a; Timsina et al., 2021). *bla*_CTX_M_ and *bla*_TEM_ were formerly detected from hospital samples (Manandhar et al., 2020)(Nayaju et al., 2021) as well as environmental samples (Thakali et al., 2020, 2021a). Similarly, *qnrS, sul1*, and *tetB* (Thakali et al., 2021a; Young et al., 2022) were detected in samples of animal and environmental samples whereas *ermB* (Timsina et al., 2021) was detected from samples taken from school children in Nepal. Other ARG that were mostly detected from clinical samples were *bla*_VEB-1_ (Tada et al., 2017), *tetA* (Nelson et al., 2020), *qnrB, dfrA, catA* and *sulII* (Manandhar et al., 2022). In Nepal, tetracycline, neomycin-doxycycline, levofloxacin, enrofloxacin, colistin, tylosin, ampicillin, amoxicillin, ceftriaxone and gentamicin are most widely prescribed and/or inappropriately used antibiotics. Likewise, chlortetracycline (CTC), bacitracin methylene disalicylate, tylosine tartarate, lincomycin, neomycin, and doxycycline are extensively used as feed additives in poultry as growth promoters (Acharya and Wilson, 2019). Over prescription and mishandling of the aforementioned drugs could have led to high presence of antibiotic resistant genes like *tetC, tet39, tetQ, tet32, tetW, tetM, tet40, tetA, tetB, aacC3, aadA, aph6, ant(3”), ant (4), ant (6), aadB, aadE, qnrB6, qnrS1, mphK, macB, macA_3, ereA2, ermCd, ermG, ermB, cepA, cfxA, bla*_TEM_, *bla*_VEB-1_, *bla*_CTX-M_, and *bla*_EC_ (Table 3, Figure 6). Among the samples, the highest ARG subtypes (n=37) were found in poultry samples (n=3) (Table 3 and Figure 5, Figure 6), which could be because of heavy antibiotics use in poultry farming in Nepal (Acharya and Wilson, 2019). A large number of ARG subtypes (n=27) were also detected in human samples, perhaps caused by over prescription, prolonged usage, self-medication practices and easy availability of antibiotics in Nepal (Dahal and Chaudhary, 2018; Acharya and Wilson, 2019). We detected ARG subtypes *tetQ* and *ermF* abundantly in human and poultry samples (Table 4 and Figure 5, Figure 6). These genes have been widely detected in human gut microbiomes (*Bacteroides fragilis* group) and in normal flora of poultry cloacae; and are found to be circulating among those microbiomes with high HRT activity (Arzese et al., 2000, Yang et al., 2014).

Our network analysis showed some strongest association between certain bacteria-*Eubacterium* and *Faecalibacterium, Eubacterium* and *Roseburia, Eubacterium* and *Collinesella, Faecalibacterium* and *Lachnospira*, and *Feacalibacterium* and *Bacteroides*. Some strongly associated AMR genes among these bacteria were *qnr, ermD, arnA*, and *aac* (Figure 7), with indication of active horizontal transfer dynamics between human, animal and environmental samples. With two nearby hospitals discharging their untreated waste into the river, our study might have picked up ARG originating from the hospitals.

**Figure 7.** Visualization of sampling site of this study i-Thapathali temporary settlement. The map was created using QGIS (an open source GIS platform) using basemaps from OpenStreetMap (25) and shape files from OpendataNepal.com (26)

Many of the ARG detected in our study are known to transcribe for the proteins that help in replication and executive functioning of bacterial cell (Supplementary Table 4) (Ochman et al., 2000). Some of the genes that code for ARG (*tetR, vanY, PBP 1A, BtgB*) were mostly associated with gut microbiome (Penders et al., 2013). In poultry samples, integrases and transposase of *IS66* and *IS21* family that help in AMR transference (Siguier et al., 2014) were also detected. Similarly, integrase and transposase of *IS116/IS110/IS902, IS30, IS605, IS200* and *IS4* family were detected in human samples that might help various bacteria to transport virulence factor, AMR genes and genes that causes overexpression of AMR genes (Bruton and Chater, 1987; Leskiw et al., 1990; Lysnyansky et al., 2009; Mohammad et al., 2020).

Five commensal enteric bacteria were predominately found in healthy human gut of Indian population (Bag et al., 2019). These bacteria were believed to act as ARG reservoir and could play a significant role in spreading ARG in enteric pathogens. Various studies also suggest gut microbiome acting as reservoir for AMR and their emergence (Forsberg et al., 2014)(Penders et al., 2013; Langelier et al., 2019; Carvalho et al., 2022). This might be of enormous epidemiological significance since novel ARG are the starting point of AMR emergence (Crofts et al., 2017). Metagenomics studies have helped us understand resistome development in gut microbiome (Miller et al., 2013; Penders et al., 2013). Finding the sources of ARG are probably the most important step in the fight against AMR.

However, working with metagenomic data is challenging and connecting each ARG to its host is still very difficult (Chiu and Miller, 2019). Thus, clinical metagenomics approach has more accuracy and will be more applicable in country like Nepal (d’Humières et al., 2021). Furthermore, sampling and preparation of DNA greatly influence findings in metagenomics analysis; improper DNA prepping might lead to insufficient genomic coverage of pathogen and ARG (Ruppé et al., 2017).

Contemporary AMR surveillance is primarily based on laboratory reporting-focusing on specific pathogens isolated only from human clinical samples (Weston et al., 2017; Hay et al., 2018). This conventional way is often lengthy, produces incomparable data and covers narrow pathogen spectrum. It is also limited and excludes AMR genes present in the commensal flora of an individual (Hendriksen et al., 2019). Increasing population density, haphazard use of antimicrobial agents, drastic changes in wildlife habitats, and increasing international travel and trade have increased the threat of AMR related infections to spread globally (Carroll et al., 2014). Deployment of active global surveillance systems for early detection of spread of zoonotic and other infectious diseases can mitigate and control outbreaks (Newell et al., 2010).

## Conclusion

Monitoring and surveillance of infectious diseases using One Health based metagenomics approach can provide important information on the emerging AMR burden. The ARG profile and antibiotic resistance (AMR) dynamics, horizontal gene transfer (HGT) events within microbiota, and virulence factor determination that indicate infection risks are some of the important information needed to truly understand overall AMR burden. Our study revealed the gut microbiome of both humans and animals can serve as a reservoir for ARG and through HGT can spillover to other environmental organisms.

## Acknowledgement

We are grateful to NHRC, Government of Nepal for providing us permission to conduct this research. We thank field team of CMDN led by Mr Bishwo P. Shrestha for their tireless work in collecting samples for this study. We are also grateful for Massachusetts Institute of Technology (MIT) for providing BioBot Automatic Sampler that was used for part of the sampling activities.

## Figures

**Figure 8.** Bioinformatics data analysis workflow used in this study.

**Figure 9.** Bar plot showing all Bacterial phylum classification in various samples according to a. 16s rRNA sequencing and b. Shotgun sequencing which was obtained from MetaPhlan V2.0. The plots were generated using ggplot2 in R studio 2022.07.1 Build 554

**Figure 10.** Prevalence of virus family in the samples obtained from Metagenomic sequencing data. a. Family wise distribution of viruses (phage) detected b. Bar plot of all the different types of phages detected in different samples, The bar plot was generated using ggplot2 in R studio 2022.07.1 Build 554 with R version 4.2.0.

**Figure 11.** Visualization of ARG profile of different samples obtained from shot-gun sequencing obtained from SHORTBred pipeline. a) Heat map clustering analysis of ARG profile of different samples with their sources, performed on ARG classified at Class level with clustering distance calculated from Bray-Curtis Index under clustering algorithm Ward. b) Alpha diversity analysis of ARG’s performed at class level (class of ARG) while measuring diversity with Shannon. The ARG diversity is significant with P-value of 0.019166 tested using ANOVA. c) Ordination analysis and significance testing for significant similarities between the different sample type and different sample sources. Non-metric multidimensional scaling (NMDS) ordination method was used with Bray-Curtis Index and statistically tested with permutational MANOVA (PERMANOVA) (F-value=2.187, R-squared=0.38456 and P-value=<0.003). Samples with statistically significant ARG association are highlighted with light green shade. d) Hierarchical clustering (dendrogram) of different samples based on the ARG profile found on them. The clustering was performed with algorithm Ward and Bray-Curtis Index as a distance measure algorithm.

**Figure 12.** Association network analysis of ARG with samples. The analysis was carried out with abundance data with associated metadata file on Gephi V0.92. The size of the circle represents number and strength of association-larger circle means a greater number of ARGs in the sample and also has more association with ARGs and other samples. The thickness of connections indicates strength of the association.

**Figure 13.** Integrative analysis of AMR gene and Taxonomic profile of different samples performed with MIC analysis for association network analysis using ResistoXplorer web platform. The color and shape of nodes are based on the type of data or profile (Resistome: yellow square; Microbiome: purple circle). In the network, the size of node represents the network centrality-based measure (degree or betweenness). While the color and width of edges shows the strength of correlation between them (MIC: varies from 0 to 1). A) Node size is based on Degree while B) Node size based on Betweenness

## Supplementary table caption

Supplementary Table 1: List of Bacterial species found in different samples via NGS sequencing

Supplementary Table 2: List of Bacteria found in different samples via 16s rRNA sequencing

Supplementary Table 3: Proteins transferred in HGT between two clades of bacteria

Supplementary Table 4: Virulence factors detected from Metagenomic sequencing data obtained from ShortBRED with their function and source (sample type).

Supplementary Table 5: Samples and its detail that was collected from Thapathali temporary settlement for this study.

