## supplementary tables for "One Health Assessment of an Urban Temporary Settlement Reveals Gut Microbiome Serving as Antimicrobial Resistance Gene Reservoir"

**Supplementary Table 1: List of Bacterial species found in different samples via NGS sequencing**

| **Bacterial species** | **Phylum** | **Sample** | **Occurrence in sample (no.)** |
| --- | --- | --- | --- |
| *Escherichia_coli* | Proteobacteria | Human and Poultry | 5 |
| *Prevotella_copri* | Bacteroidetes | Human | 10 |
| *Prevotella_sp_CAG_5226* | Bacteroidetes | Human | 1 |
| *Eubacterium_rectale* | Firmicutes | Human | 3 |
| *Lachnospira_pectinoschiza* | Firmicutes | Human | 1 |
| *Roseburia_faecis* | Firmicutes | Human | 2 |
| *Roseburia_intestinalis* | Firmicutes | Human | 1 |
| *Roseburia_inulinivorans* | Firmicutes | Human | 2 |
| *Bacteroides_plebeius* | Bacteroidetes | Human | 1 |
| *Prevotella_sp_885* | Bacteroidetes | Human | 1 |
| *Bacteroides_vulgatus* | Bacteroidetes | Human | 1 |
| *Faecalibacterium_prausnitzii* | Firmicutes | Human | 2 |
| *Prevotella_sp_CAG_520* | Bacteroidetes | Human | 1 |
| *Roseburia_faecis* | Firmicutes | Human | 2 |
| *Butyrivibrio_sp_CAG_318* | Firmicutes | Human | 1 |
| *Prevotella_sp_AM42_24* | Bacteroidetes | Human | 1 |
| *Eubacterium_siraeum* | Firmicutes | Human | 1 |
| *Chlamydia_gallinacea* | Chlamydiae | Poultry | 1 |
| *Gallibacterium_anatis* | Proteobacteria | Poultry | 1 |
| *Lawsonia_intracellularis* | Proteobacteria | Poultry | 1 |
| *Aeriscardovia_aeriphila* | Actinobacteria | Poultry | 1 |
| *Helicobacter_pullorum* | Proteobacteria | Poultry | 1 |
| *Phascolarctobacterium_sp_CAG_266* | Firmicutes | Poultry | 1 |
| *Phascolarctobacterium_sp_CAG_266* | Firmicutes | Poultry | 1 |
| *Arcobacter_cryaerophilus* | Proteobacteria | Environmental | 1 |
| *Pseudomonas aeruginosa* | Proteobacteria | Environmental | 1 |
| *Acinetobacter_johnsonii* | Proteobacteria | Environmental | 1 |
| *Acinetobacter_schindleri* | Proteobacteria | Environmental | 1 |
| *Aeromonas_caviae* | Proteobacteria | Environmental | 1 |

**Supplementary Table 2: List of Bacteria found in different samples via 16s rRNA sequencing**

| **Bacteria** | **Samples** |
| --- | --- |
| Bacteria_Actinobacteria_Actinobacteria_Bifidobacteriales_Bifidobacteriaceae_Aeriscardovia_Aeriscardovia aeriphila | Poultry |
| Bacteria_Actinobacteria_Actinobacteria_Bifidobacteriales_Bifidobacteriaceae_Bifidobacterium longum subsp. longum | Human |
| Bacteria_Actinobacteria_Actinobacteria_Corynebacteriales_Corynebacteriaceae_Corynebacterium 1_ | Poultry |
| Bacteria_Actinobacteria_Actinobacteria_Corynebacteriales_Nocardiaceae_Rhodococcus_Rhodococcus fascians | Poultry and Human |
| Bacteria_Actinobacteria_Actinobacteria_Micrococcales_Dermabacteraceae_Brachybacterium_ | Poultry |
| Bacteria_Actinobacteria_Actinobacteria_Micrococcales_Sanguibacteraceae_Sanguibacter_ | Poultry |
| Bacteria_Bacteroidetes_Bacteroidia_Bacteroidales_Bacteroidaceae_Bacteroidaceae bacterium DJFB220 | Human |
| Bacteria_Bacteroidetes_Bacteroidia_Bacteroidales_Bacteroidaceae_Bacteroides_Bacteroides caecigallinarum | Poultry |
| Bacteria_Bacteroidetes_Bacteroidia_Bacteroidales_Bacteroidaceae_Bacteroides_Bacteroides dorei | Human |
| Bacteria_Bacteroidetes_Bacteroidia_Bacteroidales_Bacteroidaceae_Bacteroides_Bacteroides fragilis | Human |
| Bacteria_Bacteroidetes_Bacteroidia_Bacteroidales_Bacteroidaceae_Bacteroides_Bacteroides massiliensis | Human |
| Bacteria_Bacteroidetes_Bacteroidia_Bacteroidales_Bacteroidaceae_Bacteroides_Bacteroides plebeius | Poultry and Human |
| Bacteria_Bacteroidetes_Bacteroidia_Bacteroidales_Bacteroidaceae_Bacteroides_Bacteroides salanitronis DSM 18170 | Poultry |
| Bacteria_Bacteroidetes_Bacteroidia_Bacteroidales_Bacteroidaceae_Bacteroides_Bacteroides sp. SB5 | Poultry |
| Bacteria_Bacteroidetes_Bacteroidia_Bacteroidales_Bacteroidaceae_Bacteroides_Bacteroides uniformis | Human |
| Bacteria_Bacteroidetes_Bacteroidia_Bacteroidales_Bacteroidaceae_Bacteroides_Mediterranea massiliensis | Poultry |
| Bacteria_Bacteroidetes_Bacteroidia_Bacteroidales_Bacteroidaceae_Bacteroides_uncultured Bacteroidales bacterium | Poultry |
| Bacteria_Bacteroidetes_Bacteroidia_Bacteroidales_Barnesiellaceae_Barnesiella_ | Human |
| Bacteria_Bacteroidetes_Bacteroidia_Bacteroidales_Marinifilaceae_Odoribacter_ | Human |
| Bacteria_Bacteroidetes_Bacteroidia_Bacteroidales_Prevotellaceae_Alloprevotella_ | Human |
| Bacteria_Bacteroidetes_Bacteroidia_Bacteroidales_Prevotellaceae_Alloprevotella_uncultured Alloprevotella sp. | Human |
| Bacteria_Bacteroidetes_Bacteroidia_Bacteroidales_Prevotellaceae_Paraprevotella_ | Poultry and Human |
| Bacteria_Bacteroidetes_Bacteroidia_Bacteroidales_Prevotellaceae_Prevotella 2_ | Human |
| Bacteria_Bacteroidetes_Bacteroidia_Bacteroidales_Prevotellaceae_Prevotella 7_Massiliprevotella massiliensis | Human |
| Bacteria_Bacteroidetes_Bacteroidia_Bacteroidales_Prevotellaceae_Prevotella 9_ | Poultry and Human |
| Bacteria_Bacteroidetes_Bacteroidia_Bacteroidales_Rikenellaceae_Alistipes_ | Human |
| Bacteria_Bacteroidetes_Bacteroidia_Bacteroidales_Rikenellaceae_Alistipes_gut metagenome | Human |
| Bacteria_Bacteroidetes_Bacteroidia_Bacteroidales_Rikenellaceae_Alistipes_uncultured Alistipes sp. | Human |
| Bacteria_Bacteroidetes_Bacteroidia_Bacteroidales_Rikenellaceae_Rikenellaceae RC9 gut group_metagenome | Human |
| Bacteria_Bacteroidetes_Bacteroidia_Bacteroidales_Tannerellaceae__ | Poultry |
| Bacteria_Bacteroidetes_Bacteroidia_Bacteroidales_Tannerellaceae_Parabacteroides_ | Poultry and Human |
| Bacteria_Bacteroidetes_Bacteroidia_Cytophagales_Cyclobacteriaceae__ | Poultry |
| Bacteria_Bacteroidetes_Bacteroidia_Sphingobacteriales_Sphingobacteriaceae_Pedobacter_ | Poultry and Human |
| Bacteria_Bacteroidetes_Bacteroidia_Sphingobacteriales_Sphingobacteriaceae_Pedobacter_Pedobacter sp. WF1 | Poultry |
| Bacteria_Bacteroidetes_Bacteroidia_Sphingobacteriales_Sphingobacteriaceae_Sphingobacterium_ | Poultry |
| Bacteria_Bacteroidetes_Bacteroidia_Sphingobacteriales_Sphingobacteriaceae_Sphingobacterium gobiense | Poultry |
| Bacteria_Bacteroidetes_Bacteroidia_Sphingobacteriales_Sphingobacteriaceae_uncultured Sphingobacterium sp. | Poultry |
| Bacteria_Bacteroidetes_Bacteroidia_Sphingobacteriales_Sphingobacteriaceae_uncultured_Sphingobacterium jejuense | Poultry |
| Bacteria_Cyanobacteria_Oxyphotobacteria_Chloroplast_Phaseolus acutifolius (tepary bean) | Poultry |
| Bacteria_Deferribacteres_Deferribacteres_Deferribacterales_Deferribacteraceae_Mucispirillum_ | Poultry |
| Bacteria_Epsilonbacteraeota_Campylobacteria_Campylobacterales_Campylobacteraceae_Campylobacter troglodytis | Human |
| Bacteria_Firmicutes_Bacilli_Bacillales_Planococcaceae_Kurthia_ | Poultry |
| Bacteria_Firmicutes_Bacilli_Bacillales_Planococcaceae_Solibacillus_ | Poultry |
| Bacteria_Firmicutes_Bacilli_Bacillales_Staphylococcaceae_Staphylococcus_ | Poultry |
| Bacteria_Firmicutes_Bacilli_Lactobacillales_Aerococcaceae_Aerococcus_ | Poultry |
| Bacteria_Firmicutes_Bacilli_Lactobacillales_Carnobacteriaceae_Alkalibacterium_ | Poultry |
| Bacteria_Firmicutes_Bacilli_Lactobacillales_Carnobacteriaceae_Desemzia_ | Poultry |
| Bacteria_Firmicutes_Bacilli_Lactobacillales_Enterococcaceae_Enterococcus_ | Poultry |
| Bacteria_Firmicutes_Bacilli_Lactobacillales_Lactobacillaceae_Lactobacillus_Lactobacillus agilis | Poultry |
| Bacteria_Firmicutes_Bacilli_Lactobacillales_Lactobacillaceae_Lactobacillus_Lactobacillus alvi | Poultry |
| Bacteria_Firmicutes_Bacilli_Lactobacillales_Lactobacillaceae_Lactobacillus_Lactobacillus aviarius | Poultry |
| Bacteria_Firmicutes_Bacilli_Lactobacillales_Lactobacillaceae_Lactobacillus_Lactobacillus delbrueckii | Human |
| Bacteria_Firmicutes_Bacilli_Lactobacillales_Lactobacillaceae_Lactobacillus_Lactobacillus johnsonii | Poultry |
| Bacteria_Firmicutes_Bacilli_Lactobacillales_Lactobacillaceae_Lactobacillus_Lactobacillus reuteri | Poultry |
| Bacteria_Firmicutes_Bacilli_Lactobacillales_Lactobacillaceae_Lactobacillus_Lactobacillus salivarius | Poultry |
| Bacteria_Firmicutes_Bacilli_Lactobacillales_Streptococcaceae_Streptococcus salivarius subsp. salivarius | Poultry |
| Bacteria_Firmicutes_Clostridia_Clostridiales_Christensenellaceae_Christensenellaceae R-7 group_ | Poultry and Human |
| Bacteria_Firmicutes_Clostridia_Clostridiales_Christensenellaceae_Christensenellaceae R-7 group_gut metagenome | Human |
| Bacteria_Firmicutes_Clostridia_Clostridiales_Clostridiaceae 1_Clostridium sensu stricto 1_Clostridium perfringens | Human |
| Bacteria_Firmicutes_Clostridia_Clostridiales_Lachnospiraceae_[Eubacterium] eligens group_ | Human |
| Bacteria_Firmicutes_Clostridia_Clostridiales_Lachnospiraceae_[Eubacterium] hallii group_ | Human |
| Bacteria_Firmicutes_Clostridia_Clostridiales_Lachnospiraceae_[Eubacterium] xylanophilum group_ | Human |
| Bacteria_Firmicutes_Clostridia_Clostridiales_Lachnospiraceae_[Ruminococcus] gauvreauii group_ | Human |
| Bacteria_Firmicutes_Clostridia_Clostridiales_Lachnospiraceae_[Ruminococcus] gnavus group_ | Human |
| Bacteria_Firmicutes_Clostridia_Clostridiales_Lachnospiraceae_[Ruminococcus] torques group_ | Poultry and Human |
| Bacteria_Firmicutes_Clostridia_Clostridiales_Lachnospiraceae_Agathobacter_ | Human |
| Bacteria_Firmicutes_Clostridia_Clostridiales_Lachnospiraceae_Anaerostipes_ | Human |
| Bacteria_Firmicutes_Clostridia_Clostridiales_Lachnospiraceae_Blautia_ | Human |
| Bacteria_Firmicutes_Clostridia_Clostridiales_Lachnospiraceae_Butyrivibrio_ | Human |
| Bacteria_Firmicutes_Clostridia_Clostridiales_Lachnospiraceae_Cellulosilyticum_ | Poultry |
| Bacteria_Firmicutes_Clostridia_Clostridiales_Lachnospiraceae_Coprococcus 1_ | Human |
| Bacteria_Firmicutes_Clostridia_Clostridiales_Lachnospiraceae_Coprococcus 2_ | Human |
| Bacteria_Firmicutes_Clostridia_Clostridiales_Lachnospiraceae_Coprococcus 3_ | Human |
| Bacteria_Firmicutes_Clostridia_Clostridiales_Lachnospiraceae_Dorea_ | Human |
| Bacteria_Firmicutes_Clostridia_Clostridiales_Lachnospiraceae_Epulopiscium_Niameybacter massiliensis | Poultry |
| Bacteria_Firmicutes_Clostridia_Clostridiales_Lachnospiraceae_Fusicatenibacter_ | Human |
| Bacteria_Firmicutes_Clostridia_Clostridiales_Lachnospiraceae_Lachnoclostridium_ | Poultry and Human |
| Bacteria_Firmicutes_Clostridia_Clostridiales_Lachnospiraceae_Lachnospira_ | Human |
| Bacteria_Firmicutes_Clostridia_Clostridiales_Lachnospiraceae_Lachnospiraceae N007 group_ | Human |
| Bacteria_Firmicutes_Clostridia_Clostridiales_Lachnospiraceae_Lachnospiraceae NK4A136 group_ | Human |
| Bacteria_Firmicutes_Clostridia_Clostridiales_Lachnospiraceae_Roseburia_ | Human |
| Bacteria_Firmicutes_Clostridia_Clostridiales_Lachnospiraceae_Tyzzerella 4_ | Human |
| Bacteria_Firmicutes_Clostridia_Clostridiales_Peptostreptococcaceae_Romboutsia_ | Poultry and Human |
| Bacteria_Firmicutes_Clostridia_Clostridiales_Peptostreptococcaceae_Terrisporobacter_ | Poultry |
| Bacteria_Firmicutes_Clostridia_Clostridiales_Ruminococcaceae_[Eubacterium] coprostanoligenes group_ | Poultry and Human |
| Bacteria_Firmicutes_Clostridia_Clostridiales_Ruminococcaceae_Butyricicoccus_ | Poultry and Human |
| Bacteria_Firmicutes_Clostridia_Clostridiales_Ruminococcaceae_Faecalibacterium_ | Human |
| Bacteria_Firmicutes_Clostridia_Clostridiales_Ruminococcaceae_Fournierella_ | Poultry and Human |
| Bacteria_Firmicutes_Clostridia_Clostridiales_Ruminococcaceae_Oscillibacter_ | Poultry and Human |
| Bacteria_Firmicutes_Clostridia_Clostridiales_Ruminococcaceae_Ruminiclostridium 6_ | Human |
| Bacteria_Firmicutes_Clostridia_Clostridiales_Ruminococcaceae_Ruminiclostridium 9_ | Poultry |
| Bacteria_Firmicutes_Clostridia_Clostridiales_Ruminococcaceae_Ruminococcaceae NK4A214 group_ | Human |
| Bacteria_Firmicutes_Clostridia_Clostridiales_Ruminococcaceae_Ruminococcaceae NK4A214 group_gut metagenome | Human |
| Bacteria_Firmicutes_Clostridia_Clostridiales_Ruminococcaceae_Ruminococcaceae UCG-002_ | Human |
| Bacteria_Firmicutes_Clostridia_Clostridiales_Ruminococcaceae_Ruminococcaceae UCG-003_ | Human |
| Bacteria_Firmicutes_Clostridia_Clostridiales_Ruminococcaceae_Ruminococcaceae UCG-005_ | Poultry and Human |
| Bacteria_Firmicutes_Clostridia_Clostridiales_Ruminococcaceae_Ruminococcaceae UCG-005_human gut metagenome | Human |
| Bacteria_Firmicutes_Clostridia_Clostridiales_Ruminococcaceae_Ruminococcaceae UCG-010_ | Human |
| Bacteria_Firmicutes_Clostridia_Clostridiales_Ruminococcaceae_Ruminococcaceae UCG-010_metagenome | Human |
| Bacteria_Firmicutes_Clostridia_Clostridiales_Ruminococcaceae_Ruminococcaceae UCG-014_ | Poultry and Human |
| Bacteria_Firmicutes_Clostridia_Clostridiales_Ruminococcaceae_Ruminococcus 1_ | Human |
| Bacteria_Firmicutes_Clostridia_Clostridiales_Ruminococcaceae_Ruminococcus 1_gut metagenome | Human |
| Bacteria_Firmicutes_Clostridia_Clostridiales_Ruminococcaceae_Subdoligranulum_ | Poultry and Human |
| Bacteria_Firmicutes_Erysipelotrichia_Erysipelotrichales_Erysipelotrichaceae_Catenibacterium_ | Human |
| Bacteria_Firmicutes_Erysipelotrichia_Erysipelotrichales_Erysipelotrichaceae_Holdemanella_ | Human |
| Bacteria_Firmicutes_Erysipelotrichia_Erysipelotrichales_Erysipelotrichaceae_Turicibacter_ | Poultry and Human |
| Bacteria_Firmicutes_Negativicutes_Selenomonadales_Acidaminococcaceae_Phascolarctobacterium_ | Poultry and Human |
| Bacteria_Firmicutes_Negativicutes_Selenomonadales_Veillonellaceae_Dialister_ | Human |
| Bacteria_Firmicutes_Negativicutes_Selenomonadales_Veillonellaceae_Megamonas_ | Human |
| Bacteria_Firmicutes_Negativicutes_Selenomonadales_Veillonellaceae_Megasphaera_ | Human |
| Bacteria_Firmicutes_Negativicutes_Selenomonadales_Veillonellaceae_Megasphaera_Megasphaera elsdenii | Human |
| Bacteria_Firmicutes_Negativicutes_Selenomonadales_Veillonellaceae_Veillonella_ | Human |
| Bacteria_Fusobacteria_Fusobacteriia_Fusobacteriales_Fusobacteriaceae_Fusobacterium_ | Poultry and Human |
| Bacteria_Proteobacteria_Alphaproteobacteria_Caulobacterales_Caulobacteraceae_Brevundimonas_ | Poultry and Human |
| Bacteria_Proteobacteria_Alphaproteobacteria_Rhizobiales_Xanthobacteraceae_Bradyrhizobium_ | Human |
| Bacteria_Proteobacteria_Alphaproteobacteria_Rhodobacterales_Rhodobacteraceae_Paracoccus_ | Poultry and Human |
| Bacteria_Proteobacteria_Alphaproteobacteria_Sneathiellales_Sneathiellaceae_Taonella_ | Poultry |
| Bacteria_Proteobacteria_Alphaproteobacteria_Sphingomonadales_Sphingomonadaceae_Sphingomonas_ | Poultry and Human |
| Bacteria_Proteobacteria_Deltaproteobacteria_Desulfovibrionales_Desulfovibrionaceae_Desulfovibrio_ | Poultry and Human |
| Bacteria_Proteobacteria_Deltaproteobacteria_Desulfovibrionales_Desulfovibrionaceae_Desulfovibrio piger | Poultry |
| Bacteria_Proteobacteria_Deltaproteobacteria_Desulfovibrionales_Desulfovibrionaceae_Lawsonia_ | Poultry |
| Bacteria_Proteobacteria_Gammaproteobacteria_Aeromonadales_Succinivibrionaceae_Succinivibrio_ | Human |
| Bacteria_Proteobacteria_Gammaproteobacteria_Betaproteobacteriales_Burkholderiaceae_Achromobacter_ | Poultry and Human |
| Bacteria_Proteobacteria_Gammaproteobacteria_Betaproteobacteriales_Burkholderiaceae_Comamonas_ | Human |
| Bacteria_Proteobacteria_Gammaproteobacteria_Betaproteobacteriales_Burkholderiaceae_Cupriavidus_ | Human |
| Bacteria_Proteobacteria_Gammaproteobacteria_Betaproteobacteriales_Burkholderiaceae_Hydrogenophaga_ | Poultry and Human |
| Bacteria_Proteobacteria_Gammaproteobacteria_Betaproteobacteriales_Burkholderiaceae_Parasutterella_ | Poultry and Human |
| Bacteria_Proteobacteria_Gammaproteobacteria_Betaproteobacteriales_Burkholderiaceae_Sutterella_ | Human |
| Bacteria_Proteobacteria_Gammaproteobacteria_Enterobacteriales_Enterobacteriaceae_Citrobacter_ | Human |
| Bacteria_Proteobacteria_Gammaproteobacteria_Enterobacteriales_Enterobacteriaceae_Enterobacter_ | Human |
| Bacteria_Proteobacteria_Gammaproteobacteria_Enterobacteriales_Enterobacteriaceae_Escherichia-Shigella_ | Poultry and Human |
| Bacteria_Proteobacteria_Gammaproteobacteria_Enterobacteriales_Enterobacteriaceae_Klebsiella_ | Human |
| Bacteria_Proteobacteria_Gammaproteobacteria_Pasteurellales_Pasteurellaceae_Gallibacterium_ | Poultry |
| Bacteria_Proteobacteria_Gammaproteobacteria_Pasteurellales_Pasteurellaceae_Gallibacterium_Gallibacterium anatis | Poultry |
| Bacteria_Proteobacteria_Gammaproteobacteria_Pasteurellales_Pasteurellaceae_Haemophilus_ | Human |
| Bacteria_Proteobacteria_Gammaproteobacteria_Pseudomonadales_Moraxellaceae_Acinetobacter_ | Poultry |
| Bacteria_Proteobacteria_Gammaproteobacteria_Pseudomonadales_Moraxellaceae_Acinetobacter_Acinetobacter lwoffii | Poultry |
| Bacteria_Proteobacteria_Gammaproteobacteria_Pseudomonadales_Pseudomonadaceae_Pseudomonas pertucinogena | Poultry |
| Bacteria_Proteobacteria_Gammaproteobacteria_Pseudomonadales_Pseudomonadaceae_Pseudomonas sp. FS29 | Poultry |
| Bacteria_Proteobacteria_Gammaproteobacteria_Xanthomonadales_Xanthomonadaceae_Lysobacter_ | Poultry |
| Bacteria_Tenericutes_Mollicutes_Anaeroplasmatales_Anaeroplasmataceae_Anaeroplasma_gut metagenome | Poultry |

**Supplementary Table 3: Proteins transferred in HGT between two clades of bacteria**

|  | **Transferrance** | |  |
| --- | --- | --- | --- |
| **Sample** | **Clade A** | **Clade B** | **Transferred Protein** |
| ES060 | Betaproteobacteria | Gammaproteobacteria | RNA-directed DNA polymerase (Reverse transcriptase) |
| ES060 | Pseudoxanthomonas_sp_GW2 | Alcanivorax_pacificus | Copper resistance protein B |
| ESD060 | Thauera_sp_27 | Dechloromonas_aromatica | HTH OST-type domain-containing protein, Integrase, & Transposase IS3/IS911 |
| ESD060 | Neisseria_meningitidis | Acinetobacter | SLT domain-containing protein |
| ESD060 | Methylomonas_methanica | Methylomicrobium_album | Elongation factor Tu 1 & 30S ribosomal protein S10 |
| ESD060 | Cupriavidus_sp_HMR_1 | Gammaproteobacteria | DDE_Tnp_IS1595 domain-containing protein |
| ESD060 | Methanoregula_boonei | Methanoregula_formicica | Methyl-coenzyme M reductase operon protein C & Methyl-coenzyme M reductase subunit gamma |
| TA330 | Bacteroides_coprophilus | Bacteroides_xylanisolvens | IS66 family element & transposase |
| TA330 | Bacillales | Clostridium_bifermentans | FMN-dependent NADH-azoreductase 3 & BFD-like [2Fe-2S] binding domain protein |
| TA330 | Lawsonia_intracellularis | Sphingobacterium_paucimobilis | 50S ribosomal protein L11 & 50S ribosomal protein L28 |
| TA340 | Histophilus_somni | Actinobacillus_pleuropneumoniae (CAUSES disease of swine DOI: https://doi.org/10.3389/fvets.2020.569370) | Relaxase |
| TA340 | Clostridium_clostridioforme | Anaerotruncus_colihominis | Transposase |
| TA340 | Gallibacterium | Actinobacillus | 1,4-alpha-glucan branching enzyme GlgB, Transposase |
| TA340 | Bacteroides_xylanisolvens | Bacteroides_finegoldii | AAA_23 domain-containing protein |
| TA340 | Prevotella_oris | Bacteroides | Reverse transcriptase domain-containing protein |
| TA340 | Prevotella | Bacteroides | Clindamycin resistance transfer factor BtgB |
| TA340 | Faecalibacterium_prausnitzii | Pseudoflavonifractor_capillosus | Mannonate dehydratase, Oxidoreductase, short chain dehydrogenase/reductase family protein |
| TA340 | Lactobacillus_salivarius | Lactobacillus_reuteri | IstB_IS21 domain-containing protein & Integrase core domain protein |
| TA340 | Oscillibacter | Ruminococcaceae | 30S ribosomal protein S12 |
| TA340 | Phascolarctobacterium_succinatutens | Virgibacillus_halodenitrificans | Cold shock protein & Peptidyl-prolyl cis/trans isomerases (PPIases) |
| TH910 | Veillonella_parvula | Veillonella_atypica | Phage portal protein, lambda family |
| TH910 | Eubacterium | Blautia | Resolvase, Protein kinase domain, Adapter protein mecA 2, Flagellin, CAAX amino terminal protease self-immunity, Peptidase_A24 domain-containing protein |
| TH910 | Bacteroides_ovatus | Bacteroides_fragilis | AAA domain protein |
| TH910 | Parabacteroides_merdae | Bacteroides | Transcriptional regulator AraC family, Type II site-specific deoxyribonuclease; FMN reductase [NAD(P)H] |
| TH910 | Ruminococcus_obeum | Ruminococcus_sp_JC304 | Shikimate dehydrogenase (NADP(+)) |
| TH910 | Eubacterium_ventriosum | Odoribacter_splanchnicus | YqeY-like protein; RNA polymerase, sigma-24 subunit, ECF subfamily |
| TH910 | Eubacterium_rectale | Roseburia_hominis | Tartrate dehydratase subunit alpha; Fumerase_C domain-containing proteiN, ABC transporter domain-containing pROTEIN; FtsX domain-containing protein; Alanyl-tRNA synthetase; Butyryl-CoA:acetate CoA-transferase; Glycosyltransferase Family 2 candid...Glycosyltransferase Family 2 candid; SAM-dependent methyltransferase: AAA domain-containing protein; Flavodoxin-like domain-containing protein; NADH-dependent butanol; RNA polymerase sigma factor, sigma-70 familydehydrogenasw and Non-specific serine/threonine protein kinase; |
| TH910 | Faecalibacterium_prausnitzii | Subdoligranulum_sp_4_3_54A2FAA | ABC-type multidrug transport system, ATPase and permease components |
| TH910 | Ruminococcus_albus | Faecalibacterium_prausnitzii | Putative 23S rRNA m2A2503 methyltransferase; Histidinol-phosphatase |
| TH910 | Eubacterium_rectale | Ruminococcus_obeum | Integrase core domain protein |
| TH910 | Ruminococcaceae | Clostridium | Bacterial mobilization protein MobC; Transcriptional regulator, AraC family |
| TH910 | Bacteroides_massiliensis | Bacteroides_vulgatus | SIR2_2 domain-containing protein; HTH araC/xylS-type domain-containing protein, |
| TH910 | Dorea_longicatena | Coprococcus_comes | V-type ATP synthase subunit I |
| TH910 | Alistipes | Bacteroides | DAK2 domain fusion protein YloV |
| TH910 | Subdoligranulum_variabile | Faecalibacterium_prausnitzii | DAK2 domain fusion protein YloV |
| TH910 | Eubacterium_rectale | Butyrivibrio_crossotus | NAD dependent epimerase/dehydratase |
| TH910 | Faecalibacterium_prausnitzii | Ruminococcus_sp_JC304 | Ferritin |
| TH910 | Bacteroides_clarus | Bacteroides_stercoris | Glycosyl hydrolase family 88 |
| TH910 | Firmicutes_bacterium_M10_2 | Eubacterium_biforme | AB hydrolase-1 domain-containing protein |
| TH910 | Clostridium_clostridioforme | Coprococcus_catus | Phage DNA replication protein |
| TH910 | Ruminococcus_sp_5_1_39BFAA | Eubacterium_eligens | HTH marR-type domain-containing protein;Multidrug resistance protein MATE family |
| TH910 | Faecalibacterium_prausnitzii | Roseburia | Putative efflux protein, MATE family; DNA-binding helix-turn-helix protein |
| TH910 | Clostridium_scindens | Coprococcus_eutactus | AlwI restriction endonuclease |
| TH910 | Dorea_formicigenerans | Ruminococcus_sp_5_1_39BFAA | Stage IV sporulation protein B |
| TH910 | Bacteroides_dorei | Bacteroides_sp_3_1_33FAA | SGNH_hydro domain-containing protein |
| TH940 | Roseburia_intestinalis | Lachnospiraceae_bacterium_1_4_56FAA | Lysine--tRNA ligase; Group II intron, maturase-specific domain protein |
| TH940 | Ruminococcus_sp_JC304 | Ruminococcus_obeum | Helix-turn-helix |
| TH940 | Bacteroides_plebeius | Bacteroides_coprophilus | Acyl_transf_3 domain-containing protein; Cyclic nucleotide-binding domain-containing protein |
| TH940 | Eubacterium_rectale | Butyrivibrio_crossotus | ASCH domain-containing protein; NAD dependent epimerase/dehydratase family protein |
| TH940 | Ruminococcaceae | Clostridium | AraC-type DNA-binding domain-containing proteins; |
| TH940 | Roseburia_inulinivorans | Faecalibacterium_prausnitzii | Acetyltransferases; Threonine dehydrogenase and related Zn-dependent dehydrogenases |
| TH940 | Eubacterium_rectale | Roseburia_intestinalis | Very short patch repair endonuclease; Site-specific recombinases, DNA invertase Pin homologs |
| TH940 | Roseburia_inulinivorans | Eubacterium_rectale | DUF1016_N domain-containing protein |
| TH940 | Bacteroides_sp_2_1_7 | Cronobacter_universalis | DUF6531 domain-containing protein |
| TH940 | Eubacterium_rectale | Roseburia_intestinalis | Relaxase/Mobilisation nuclease domain |
| TH940 | Bacteroidaceae | Clostridiales_noname | Conserved protein found in conjugate transposon; Homologues of TraJ from Bacteroides conjugative transposon |
| TH940 | Dorea_longicatena | Roseburia_hominis | Glycerate kinase; Ribonuclease 3 |
| TH940 | Bilophila | Bacteroidales | Efflux transporter, RND family, MFP subunit |
| TH950 | Eubacterium_eligens | Bacteroides_pectinophilus | Gluconate 5-dehydrogenase; 4-deoxy-L-threo-5-hexosulose-uronate ketol-isomerase |
| TH950 | Roseburia_intestinalis | Roseburia_inulinivorans | Site-specific recombinases, DNA invertase Pin homologs; Stage 0 sporulation protein A homolog |
| TH950 | Blautia_sp_KLE_1732 | Roseburia_intestinalis | Divergent AAA domain protein; DNA-directed RNA polymerase specialized sigma subunit, sigma24 homolog |
| TH950 | Bacteroides_pectinophilus | Eubacterium_eligens | Glyceraldehyde-3-phosphate dehydrogenase; 2,3-bisphosphoglycerate-independent phosphoglycerate mutase |
| TH950 | Bacteroides_dorei | Parabacteroides_johnsonii | Arm-DNA-bind_5 domain-containing protein |
| TH950 | Pseudoflavonifractor_capillosus | Erysipelotrichaceae_bacterium_6_1_45 | Ankyrin repeat protein |
| TH950 | Clostridium | Roseburia | SGNH_hydro domain-containing protein |
| TH950 | Bacteroides_sp_4_1_36 | Bacteroides_xylanisolvens | Site-specific DNA-methyltransferase (adenine-specific) |
| TH950 | Roseburia_intestinalis | Eubacterium_ramulus | Transposase IS4 family protein |
| TH950 | Coprococcus_comes | Lactobacillales | Glycosyltransferase, group 1 family protein |
| TH950 | Coprobacillus_sp_D7 | Eubacterium_rectale | Predicted amidohydrolase |
| TH950 | Eubacterium_eligens | Eubacterium_ventriosum | Coenzyme A biosynthesis bifunctional protein CoaBC; Aspartate 1-decarboxylase |
| TH950 | Roseburia_inulinivorans | Eubacterium_rectale | Histidinol dehydrogenase; Transposase IS116/IS110/IS902 family./Transposase |
| TH950 | Eubacterium_rectale | Coprococcus_catus | Predicted ATPase (AAA+ superfamily); Nitroreductase domain-containing protein; Adenylate kinase and related kinases |
| TH950 | Eubacterium_eligens | Clostridium_sp_L2_50 | HDc domain-containing protein; Nudix hydrolase domain-containing protein; Transposase, IS200 family |
| TH950 | Roseburia_intestinalis | Roseburia_inulinivorans | Transposase IS116/IS110/IS902 family |
| TH950 | Oribacterium_sp_oral_taxon_078 | Roseburia_intestinalis | Antitoxin; DUF2185 domain-containing protein |
| TH950 | Eubacterium_ventriosum | Lachnospiraceae | Virulence protein |
| TH950 | Lachnospiraceae | Eubacterium | Integrase; Methyltransf_11 domain-containing protein |
| TH950 | Ruminococcus_sp_JC304 | Faecalibacterium_prausnitzii | Ferritin |
| TH950 | Eubacterium_eligens | Lachnospiraceae_bacterium_6_1_63FAA | GMP synthase [glutamine-hydrolyzing]; Tyr recombinase domain-containing protein |
| TH980 | Prevotella_copri | Prevotella_stercorea | Methyltransferase domain protein; Sigma-70 region 2 |
| TH980 | Prevotella_dentalis | Prevotella_copri | ABC superfamily ATP binding cassette transporter |
| TH980 | Prevotella_sp_C561 | Prevotella_copri | Leucine-rich repeat domain-containing protein |
| TH980 | Catenibacterium_mitsuokai | Clostridiales | ABC superfamily ATP binding cassette transporter; Transposase, IS605 family |
| TH980 | Coprobacillus | Catenibacterium_mitsuokai | Transposase; Permease IIC component |
| TH980 | Prevotella_copri | Bacteroides_thetaiotaomicron | ATP-binding protein |
| TH980 | Prevotella_paludivivens | Prevotella_copri | CTP synthase; Membrane protein insertase YidC |
| TH980 | Bacteroides_vulgatus | Prevotella_copri | HAD hydrolase, family IA, variant 3 |
| TH1010 | Lachnospiraceae_bacterium_9_1_43BFAA | Eubacterium_rectale | Site-specific DNA-methyltransferase (adenine-specific); RNA methyltransferase |
| TH1010 | Clostridium_sp_M62_1 | Ruminococcus_torques | Glucosamine-6-phosphate deaminase |
| TH1010 | Butyrivibrio_crossotus | Eubacterium_ramulus | NAD dependent epimerase/dehydratase family protein; |
| TH1010 | Parabacteroides_sp_D13 | Prevotella_copri | HTH cro/C1-type domain-containing protein; TPR domain protein; Cys/Met metabolism PLP-dependent enzyme |
| TH1010 | Roseburia_inulinivorans | Eubacterium_rectale | Multidrug resistance protein, MATE family |
| TH1010 | Prevotella_copri | Prevotella_stercorea | OMP_b-brl_3 domain-containing protein; HTH luxR-type domain-containing protein; |
| TH1010 | Bacteroides | Prevotella_stercorea | Transposase, IS116/IS110/IS902 family; Transposase, IS116/IS110/IS902 family; |
| TH1010 | Clostridium_clostridioforme | Coprococcus_sp_ART55_1 | Two-component system response regulator; Predicted transcriptional regulators |
| TH1010 | Coprococcus_comes | Dorea_longicatena | Peripla_BP_4 domain-containing protein; Branched-chain amino acid ABC transporter permease protein |
| TH1010 | Eubacterium_rectale | Clostridium_bolteae | Heat-inducible transcription repressor HrcA |
| TH1020 | Sutterella_wadsworthensis | Neisseria_meningitidis | HTH gntR-type domain-containing protein |
| TH1020 | Butyrivibrio_crossotus | Ruminococcus_obeum | Replication initiator protein A domain protein; ParB-like protein; CobQ/CobB/MinD/ParA nucleotide binding domain protein |
| TH1020 | Prevotella_copri | Prevotella_bivia | DNA mismatch repair protein MutS; ATPase |
| TH1020 | Eubacterium | Ruminococcus_lactaris | Transposase and inactivated derivatives; ATP-dependent 6-phosphofructokinase |
| TH1020 | Prevotella_copri | Prevotella_stercorea | Transcription termination/antitermination factor NusG; dTDP-4-dehydrorhamnose 3,5-epimerase; |
| TH1020 | Lachnospiraceae | Eubacteriaceae | NAD dependent epimerase/dehydratase family protein; ASCH domain-containing protein; Hydrolase_4 domain-containing protein |
| TH1020 | Eubacterium_rectale | Blautia_producta | ABC superfamily ATP binding cassette transporter |
| TH1020 | Prevotella_bergensis | Parabacteroides_sp_D13 | T2SS-T3SS_pil_N domain-containing protein |
| TH1020 | Eubacterium_rectale | Megamonas | Elongation factor Ts |
| TH1020 | Megamonas_rupellensis | Megamonas_funiformis | IS605 OrfB family transposase |
| TH1020 | Ruminococcus_albus | Ruminococcus_flavefaciens | Putative 23S rRNA m2A2503 methyltransferase; |
| TH1020 | Bacteroides_sp_2_2_4 | Prevotella_stercorea | Site-specific recombinase, phage integrase family; Glycosyltransferase, WecB/TagA/CpsF family; BIG2 domain-containing protein |
| TH1020 | Prevotella_timonensis | Prevotella_stercorea | *KilA-N domain-containing protei;* Flavodoxin |
| TH1020 | Eubacterium_rectale | Lachnospiraceae | Methyl-accepting chemotaxis protein; Predicted membrane protein |
| TH1020 | Prevotella_bergensis | Prevotella_copri | DUF4377 domain-containing protein; OMP_b-brl_3 domain-containing protein |
| TH1020 | Bacteroides_thetaiotaomicron | Prevotella_copri | ATPase_2 domain-containing protein; RNA methylase SpoU family |
| TH1020 | Prevotella_stercorea | Prevotella_copri | ATP-binding protein; DUF4230 domain-containing protein |
| TH1020 | Prevotella_stercorea | Prevotella_dentalis | ABC superfamily ATP binding cassette transporter; |
| TH1030 | Dorea_longicatena | Coprococcus_catus | Glycerate kinase; HTH cro/C1-type domain-containing protein; Transcriptional regulator, MerR family; Stage 0 sporulation protein A homolog; Histidine kinase |
| TH1030 | Faecalibacterium_prausnitzii | Eubacterium_rectale | Glycine--tRNA ligase; DUF4366 domain-containing protein |
| TH1030 | Prevotella_copri | Bacteroides_thetaiotaomicron | ATP-binding protein; |
| TH1030 | Faecalibacterium_prausnitzii | Ruminococcus_albus | Histidinol-phosphatase; Putative 23S rRNA m2A2503 methyltransferase |
| TH1030 | Eubacterium_siraeum | Eubacterium_rectale | 3-dehydroquinate dehydratase; |
| TH1030 | Ruminococcus_sp_JC304 | Faecalibacterium_prausnitzii | Methylase involved in ubiquinone/menaquinone biosynthesis |
| TH1030 | Dorea_formicigenerans | Butyrivibrio_crossotus | ORF6N domain-containing protein |
| TH1030 | Clostridium_sp_ATCC_BAA_442 | Ruminococcus_sp_JC304 | Metallo-beta-lactamase domain protein |
| TH1030 | Butyricicoccus_pullicaecorum | Pseudoflavonifractor_capillosus | DUF6017 domain-containing protein |
| TH1030 | Ruminococcaceae | Lachnospiraceae | HTH_38 domain-containing protein |
| TH1030 | Lachnospiraceae | Eubacterium_rectale | Lysine decarboxylase |
| TH1030 | Prevotella_copri | Parabacteroides_sp_D13 | Argininosuccinate lyase; Orotate phosphoribosyltransferase; Plasmid recombination enzyme; SF3 helicase domain-containing protein |
| TH1030 | Bulleidia_extructa | Lachnoanaerobaculum_saburreum | Type I restriction modification DNA specificity domain protein; Site-specific recombinase, phage integrase family |
| TH1030 | Eubacterium_rectale | Ruminococcus_lactaris | Predicted Fe-S oxidoreductases; |
| TH1030 | Prevotella_stercorea | Prevotella_copri | Integral membrane protein TerC family; PepSY domain protein |
| TH1030 | Bilophila | Desulfovibrio_piger | YscQ/HrcQ family type III secretion apparatus protein; Type III secretion apparatus protein, YscR/HrcR family |
| TH1030 | Roseburia_intestinalis | Roseburia_inulinivorans | Site-specific recombinases, DNA invertase Pin homologs; Divergent AAA domain protein |
| TH1030 | Bacteroides_faecis | Prevotella_copri | Uncultured bacterium extrachromosomal DNA RGI01876 |
| TH1030 | Faecalibacterium_prausnitzii | Eubacterium_rectale | Predicted exporters of the RND superfamily; Trypsin-like serine proteases, typically periplasmic, contain C-terminal PDZ domain |
| TH1030 | Ruminococcus_sp_5_1_39BFAA | Blautia_sp_KLE_1732 | HTH domain protein |
| TH1030 | Eubacterium_rectale | Coprococcus_eutactus | RNA polymerase sigma-70 factor, ECF subfamily |
| TH1030 | Prevotella_multisaccharivorax | Prevotella_bryantii | Radical SAM domain protein; TonB-dependent receptor plug |
| TH1030 | Bacteroides_plebeius | Prevotella_copri | DUF1273 family protein |
| TH1030 | Roseburia_inulinivorans | Roseburia_hominis | Acetyltransferase, GNAT family; Transcriptional regulator |
| TH1030 | Faecalibacterium_prausnitzii | Oscillibacter | Ion-translocating oxidoreductase complex subunit B; Electron transport complex, RnfABCDGE type, A subunit; |
| TH1030 | Bacteroides_sp_2_2_4 | Bacteroides_ovatus | DNA-binding helix-turn-helix protein; |
| TH1030 | Prevotella | Parabacteroides_sp_D13 | Site-specific recombinase, phage integrase family |
| TH1030 | Roseburia_intestinalis | Dorea_formicigenerans | Transposase and inactivated derivatives, IS30 family; ATPase_AAA_core domain-containing protein |
| TH1030 | Ruminococcus_sp_JC304 | Faecalibacterium_prausnitzii | Ferritin |
| TH1030 | Roseburia_inulinivorans | Roseburia_hominis | Uridine phosphorylase; PUA domain-containing protein; Nicotinate phosphoribosyltransferase |
| TH1030 | Roseburia_inulinivorans | Bacteroides_pectinophilus | Queuine tRNA-ribosyltransferase; Chorismate synthase; ABC transporter domain-containing protein; ABC transmembrane type-1 domain-containing protein |
| TH1030 | Roseburia_inulinivorans | Coprococcus_comes | Tnp_DDE_dom domain-containing protein; DNA-binding helix-turn-helix protein |
| TH1030 | Eubacterium_ramulus | Eubacterium_rectale | ATPase |
| TH1040 | Faecalibacterium_prausnitzii | Ruminococcus_sp_JC304 | Methylase involved in ubiquinone/menaquinone biosynthesis; |
| TH1040 | Bacteroides_faecis | Bacteroides_coprocola | Plasmid recombination enzyme; OmpA-like domain-containing protein; DNA mismatch repair protein mutS; PPM-type phosphatase domain-containing protein; bPH_2 domain-containing protein; Peptidase_M23 domain-containing protein; Transposase |
| TH1040 | Roseburia_inulinivorans | Lactobacillus_ruminis | Transposase |
| TH1040 | Prevotella | Bacteroides | Pyridoxamine 5'-phosphate oxidase family protein; Putative organophosphate reductase; DUF3791 domain-containing protein; Aldo/keto reductase; DUF1273 family protein |
| TH1040 | Lactobacillales | Coprococcus_comes | Glycosyltransferase, group 1 family protein; |
| TH1040 | Prevotella_stercorea | Prevotella_copri | YccF domain-containing protein |
| TH1040 | Faecalibacterium_prausnitzii | Subdoligranulum_sp_4_3_54A2FAA | ABC-type multidrug transport system, ATPase and permease components |
| TH1040 | Prevotella_stercorea | Prevotella_copri | N-acetylmuramoyl-L-alanine amidase |
| TH1040 | Prevotella | Parabacteroides | DUF5110 domain-containing protein; Sigma-70 region 2; Methyltransferase domain protein; Plasmid recombination enzyme |
| TH1040 | Clostridium_sp_L2_50 | Faecalibacterium_prausnitzii | NUDIX hydrolase; Crossover junction endodeoxyribonuclease RuvC |
| TH1040 | Eubacterium_rectale | Roseburia_inulinivorans | TsaA-like domain-containing protein; Low specificity L-threonine aldolase |
| TH1040 | Eubacterium_rectale | Lachnospiraceae | DEDD_Tnp_IS110 domain-containing protein |
| TH1040 | Bacteroides | Prevotella_multisaccharivorax | DNA binding domain, excisionase family |
| TH1040 | Ruminococcus_albus | Faecalibacterium_prausnitzii | Histidinol-phosphatase; Putative 23S rRNA m2A2503 methyltransferase; Site-specific recombinases, DNA invertase Pin homologs |
| TH1040 | Prevotella | Roseburia_inulinivorans | ABC superfamily ATP binding cassette transporter, ABC protein; |
| TH1110 | Clostridium_nexile | Eubacterium_rectale | ABC-type sugar/spermidine/putrescine/iron/thiamine transport system, ATPase component |
| TH1110 | Roseburia_inulinivorans | Oribacterium_sp_oral_taxon_078 | Ribosomal RNA small subunit methyltransferase H; Antitoxin |
| TH1110 | Clostridiales | Actinobacteria | Transposase-like protein |
| TH1110 | Ruminococcus_sp_5_1_39BFAA | Clostridium_butyricum | Mutator family transposase |
| TH1110 | Catenibacterium_mitsuokai | Eubacterium_biforme | HTH rpiR-type domain-containing protein; Transposase, IS116/IS110/IS902 family |
| TH1110 | Prevotella_stercorea | Bacteroides | CRISPR-associated endonuclease Cas9; ATP-dependent DNA helicase, RecQ family |
| TH1110 | Parabacteroides | Prevotella | HTH cro/C1-type domain-containing proteiN:ABC superfamily ATP binding cassette transporter: Cobyrinic acid a,c-diamide synthase: Putative phosphohydrolase; Single-stranded DNA-binding protein : ABC superfamily ATP binding cassette transporter |
| TH1110 | Roseburia_inulinivorans | Lachnospiraceae_bacterium_3_1_46FAA | HTH cro/C1-type domain-containing protein |
| TH1110 | Ruminococcus_sp | Roseburia_inulinivorans | Integrase core domain protein |
| TH1110 | Eubacterium | Roseburia | RNA-directed RNA polymerase; RNA_pol_A_CTD domain-containing protein |
| TH1110 | Bacteroides | Prevotella | Site-specific recombinase phage integrase family |
| TH1110 | Lachnospiraceae | Eubacteriaceae | Adenylate kinase and related kinases; Predicted ATPase (AAA+ superfamily); Predicted ATPase (AAA+ superfamily) |
| TH1110 | Bacteroides_ovatus | Parabacteroides_sp_D13 | Peptidase A2 domain-containing protein; |
| TH1110 | Prevotella_copri | Prevotella_stercorea | DNA_pol3_alpha domain-containing protein |
| TH1110 | Bacteroides_fragilis | Prevotella_multisaccharivorax | Abortive infection protein (10.1093/nar/gkt1419) |
| TH1110 | Prevotella_copri | Prevotella_stercorea | Transcriptional regulator; Phosphatidylglycerophosphatase A; CDP-alcohol phosphatidyltransferase family protein; Inositol phosphorylceramide synthase; GtrA domain-containing protein; Transcriptional regulator LacI family; ATPase/histidine kinase/DNA gyrase B/HSP90 domain protein; Glycosyl hydrolase family 31; DUF4982 domain-containing protein; Acetyltransferase, GNAT family; Amino acid carrier protein; DUF4230 domain-containing protein; ATP-binding protein; OmpA family protein |
| TH1110 | Roseburia_inulinivorans | Roseburia_intestinalis | Fn3_like domain-containing protein; Fe2+ transport system protein A |
| TH1110 | Prevotella_copri | Riemerella_columbina | Uncultured bacterium extrachromosomal DNA RGI01876; Truncated RteA |
| TH1110 | Clostridium_difficile | Roseburia_intestinalis | Alpha-L-arabinofuranosidase; |
| TH1110 | Veillonella_sp_HPA0037 | Megasphaera_elsdenii | Transposase IS200-family protein; TetR protein |
| TH1110 | Clostridium_nexile | Eubacterium_rectale | PDZ domain-containing protein; Uncharacterized FAD-dependent dehydrogenase |
| TH1110 | Prevotella_copri | Prevotella_stercorea | AAA-ATPase_like domain-containing protein; |
| TH1110 | Roseburia_inulinivorans | Roseburia_intestinalis | Aldo_ket_red domain-containing protein; HATPase_c_5 domain-containing protein; Transposase; Integrase core domain protein; Toxin-antitoxin system antitoxin component Xre family |
| TH1120 | Ruminococcus_bromii | Butyrivibrio_crossotus | Penicillin-binding protein 1A |
| TH1120 | Prevotella_stercorea | Prevotella_copri | Dihydrofolate reductase; Glycosyl transferase group 1 family |
| TH1120 | Butyrivibrio_crossotus | Prevotella_copri | Aminoglycoside phosphotransferase; RNA polymerase sigma factor sigma-70 family |
| TH1120 | Roseburia_intestinalis | Butyrivibrio_crossotus | Transcriptional regulator, AraC family; Methyl-accepting chemotaxis protein signaling domain protein |
| TH1120 | Prevotella_copri | Prevotella_stercorea | Methyl-accepting chemotaxis protein signaling domain protein; dTDP-4-dehydrorhamnose 3,5-epimerase |
| TH1120 | Oscillibacter_valericigenes | Faecalibacterium_prausnitzii | 30S ribosomal protein S10; 50S ribosomal protein L3 |
| TH1130 | Faecalibacterium_prausnitzii | Ruminococcaceae | Histidinol-phosphatase; Putative 23S rRNA m2A2503 methyltransferase |
| TH1130 | Prevotella_stercorea | Prevotella | Transcriptional regulator; Phosphatidylglycerophosphatase A; CDP-alcohol phosphatidyltransferase family protein; Inositol phosphorylceramide synthase; GtrA domain-containing protein; Transcriptional regulator LacI family; ATPase/histidine kinase/DNA gyrase B/HSP90 domain protein; Glycosyl hydrolase family 31; DUF4982 domain-containing protein; Acetyltransferase, GNAT family; Amino acid carrier protein; DUF4230 domain-containing protein; FimB/Mfa2 family fimbrial subunit; Transcriptional regulator AraC family |
| TH1130 | Dorea_formicigenerans | Clostridiales | Transcriptional regulator |
| TH1130 | Lachnospiraceae_bacterium_7_1_58FAA | Clostridiales | Integrase catalytic domain-containing protein; |
| TH1130 | Streptococcus_salivarius | Lactobacillales | Integral membrane protein; Helper of Tim protein 13 |
| TH1130 | Blautia_sp_KLE_1732 | Firmicutes | Restriction endonuclease |
| TH1130 | Prevotella_timonensis | Prevotella | Cysteine-rich domain protein; LUD_dom domain-containing protein; Putative iron-sulfur cluster-binding protein |
| TH1130 | Prevotella_copri | Prevotella | YccF domain-containing protein |
| TH1130 | Clostridium_clostridioforme | Clostridiales | DNA topoisomerase |
| TH1130 | Eubacterium_ventriosum | Clostridiales | Predicted dehydrogenases and related proteins |
| TH1130 | Erysipelotrichales | Firmicutes | TraG family protein; ATPases involved in chromosome partitioning; Replication initiator protein A domain protein; ParB-like partition proteins; Reverse transcriptase (RNA-dependent DNA polymerase) |
| TH1130 | Prevotella_stercorea | Prevotella | Alpha amylase, catalytic domain protein; Fic family protein |
| TH1130 | Prevotella | Bacteroidales | Aldo/keto reductase; DUF3791 domain-containing protein; Putative organophosphate reductase |
| TH1130 | Bacteroides_ovatus | Bacteroidales | TonB-dependent receptor; Coproporphyrinogen-III oxidase; Transporter, major facilitator family protein |
| TH1130 | Eubacterium_rectale | Clostridiales | HPr kinase/phosphorylase; M18 family aminopeptidase |
| TH1130 | Dorea_formicigenerans | Clostridiales | Mutator family transposase |
| TH1130 | Eubacterium_rectale | Clostridiales | Transcriptional regulator |
| TH1130 | Prevotella_dentalis | Prevotella | HAD hydrolase, family IA, variant 3; ABC superfamily ATP binding cassette transporter; ATPase |
| TH1130 | Roseburia_inulinivorans | Lachnospiraceae | Arginase family; Peptide deformylase; Transcriptional regulator, Spx/MgsR family |
| TH1130 | Dorea_formicigenerans | Clostridiales | HAD superfamily hydrolase; Putative adenosine deaminase; PadR domain-containing protein; Acetyltransferases, including N-acetylases of ribosomal proteins |
| TH1130 | Lachnospiraceae | Clostridiales | RNA methyltransferase; |
| TH1130 | Prevotella_bivia | Prevotella | Anthranilate synthase |
| TH1130 | Bacteroides_stercoris | Bacteroides | MFS domain-containing protein; |
| TH1130 | Eubacterium_rectale | Clostridiales | Glycosidases |
| TH1130 | Bacteroides_sp_D1 | Bacteroidales | 30S ribosomal protein S6; TonB-dependent receptor; Lipocalin-like domain-containing protein; MotA/TolQ/ExbB proton channel family protein; CobN/magnesium chelatase family protein; ATP-dependent DNA helicase, RecQ family; DUF2023 domain-containing protein; Flavodoxin; Exonuclease; ADP-ribosylglycohydrolase; VirE N-terminal domain protein; Na+/H+ antiporter |
| TH1130 | Anaerostipes_hadrus | Clostridiales | Helix-turn-helix; Transcriptional regulator, AraC family; Cobalt transporter; ABC transporter; ABC-type multidrug transport system, ATPase and permease components |
| TH1130 | Ruminococcus | Clostridiales | VanY domain-containing protein; RNA polymerase sigma factor, sigma-70 family; ABC-type multidrug transport system, ATPase component |
| TH1130 | Roseburia_intestinalis | Clostridiales | Phosphate transporter ATP-binding protein |
| TH1130 | Parabacteroides | Bacteroidales | Receptor mostly Fe transport; Peptidase, U32 family; Homoserine O-acetyltransferase; tRNA(Ile)-lysidine synthase; 3-deoxy-manno-octulosonate cytidylyltransferase; RNA polymerase sigma-70 factor; Glycosyl hydrolase family protein; DNA helicase; DnaD domain protein; RNA polymerase sigma-43 factor; Arylsulfatase; AraC family transcriptional regulator; PAP2 family protein; Glutamine--tRNA ligase; DedA family protein; Carbohydrate kinase; Aspartate--tRNA ligase; RNA polymerase sigma factor; UDP-glucose 4-epimerase; Sigma-70 region 2; Methyltransferase domain protein; Plasmid recombination enzyme |
| TH1130 | Coprobacillus_sp_8_2_54BFAA | Firmicutes | PLDc_N domain-containing protein; |
| TH1130 | Coprobacillus_sp_3_3_56FAA | Firmicutes | HTH cro/C1-type domain-containing protein |
| TH1150 | Prevotella_stercorea | Prevotella_copri | Transcriptional regulator |

**Supplementary Table 4: Virulence factors detected from Metagenomic sequencing data obtained from ShortBRED with their function and source (sample type).**

| **VFDB database** |  | **Virulence gene** | **Occurrence in sample** | **Bacterial species source (according to VFDB)** | **Description** | **Function** |
| --- | --- | --- | --- | --- | --- | --- |
| VFG0462 |  | *csgG* | Poultry and Human | *Salmonella enterica* (serovar typhimurium) LT2 | Putative transcriptional regulator | Adherence |
| VFG0871 |  | *fimB* | Poultry and Human | *Escherichia coli*  CFT073 | Type 1 fimbriae Regulatory protein fimB | Adherence |
| VFG0872 |  | *fimE* | Poultry and Human | *Escherichia coli*  CFT073 | Type 1 fimbriae Regulatory protein fimE | Adherence |
| VFG0874 |  | *fimI* | Poultry and Human | *Escherichia coli*  CFT073 | Fimbrin-like protein fimI precursor | Adherence |
| VFG0875 |  | *fimC* | Poultry and Human | *Escherichia coli*  CFT073 | Chaperone protein fimC precursor | Adherence |
| VFG0876 |  | *fimD* | Poultry and Human | *Escherichia coli*  CFT073 | Outer membrane usher protein fimD precursor | Adherence |
| VFG0877 |  | *fimF* | Poultry | *Escherichia coli*  CFT073 | FimF protein precursor | Adherence |
| VFG0878 |  | *fimG* | Poultry | *Escherichia coli*  CFT073 | FimG protein precursor | Adherence |
| VFG0879 |  | *fimH* | Poultry and Human | *Escherichia coli*  CFT073 | FimH protein precursor | Adherence |
| VFG1223 |  | *pilT* | Poultry and environmental | *Pseudomonas aeruginosa* PAO1 | twitching motility protein PilT | Adherence |
| VFG1440 |  | *ibeC* | Human | *E. coli* | Membrane protein YijP | Invasion |
| VFG1443 |  | *ompA* | Poultry and Human | *E. coli* | Outer membrane protein A | Invasion |
| VFG1444 |  | *aslA* | Human | *E. coli* | Putative arylsulfatase | Invasion |
| VFG1445 |  | *traJ* | Poultry and Human | [Escherichia coli (strain K12)](https://www.uniprot.org/taxonomy/83333) | Regulator in cojugation | Invasion |
| VFG1448 |  | *kpsD* | Human | *E. coli* | Polysialic acid transport protein | Invasion |
| VFG1450 |  | *kpsM* | Human | *E. coli* | Polysialic acid transport protein | Invasion |
| VFG0358 |  | *psn* | Human | *Yersinia pestis* CO92 | Pesticin/yersiniabactin receptor protein | Iron uptake |
| VFG0360 |  | *ybtT* | Human | *Yersinia pestis* CO92 | Yersiniabactin biosynthetic protein YbtT | Iron uptake |
| VFG0361 |  | *ybtU* | Poultry and Human | *Yersinia pestis* CO92 | Yersiniabactin biosynthetic protein YbtU | Iron uptake |
| VFG0362 |  | *irp1* | Human | *Yersinia pestis* CO92 | Yersiniabactin biosynthetic protein HMWP1 (high molecular weight protein 1) | Iron uptake |
| VFG0363 |  | *irp2* | Poultry and Human | *Yersinia pestis* CO92 | Yersiniabactin biosynthetic protein HMWP2 (high molecular weight protein 2) | Iron uptake |
| VFG0364 |  | *ybtA* | Human | *Yersinia pestis* CO92 | Transcriptional regulator YbtA | Iron uptake |
| VFG0365 |  | *ybtP* | Human | *Yersinia pestis* CO92 | Lipoprotein inner membrane ABC-transporter | Iron uptake |
| VFG0366 |  | *ybtQ* | Poultry and Human | *Yersinia pestis* CO92 | Inner membrane ABC-transporter YbtQ | Iron uptake |
| VFG0367 |  | *ybtX* | Poultry and Human | *Yersinia pestis* CO92 | Putative signal transducer | Iron uptake |
| VFG0916 |  | *chuS* | Poultry and Human | *Escherichia coli*  CFT073 | Putative heme/hemoglobin transport protein | Iron uptake |
| VFG0917 |  | *chuA* | Human | *Escherichia coli*  CFT073 | Outer membrane heme/hemoglobin receptor | Iron uptake |
| VFG0918 |  | *chuT* | Poultry and Human | *Escherichia coli*  CFT073 | Putative Periplasmic binding protein | Iron uptake |
| VFG0919 |  | *chuW* | Human | *Escherichia coli*  CFT073 | Putative oxygen independent coproporphyrinogen III oxidase | Iron uptake |
| VFG0922 |  | *chuU* | Human | *Escherichia coli*  CFT073 | Putative permease of iron compound ABC transport system | Iron uptake |
| VFG0923 |  | *fepA* | Poultry and Human | *Escherichia coli*  CFT073 | Ferrienterobactin receptor precursor | Iron uptake |
| VFG0925 |  | *fepC* | Poultry and Human | *Escherichia coli*  CFT073 | Ferric enterobactin transport ATP-binding protein fepC | Iron uptake |
| VFG0927 |  | *fepE* | Human | *Escherichia coli*  CFT073 | Ferric enterobactin transport protein fepE | Iron uptake |
| VFG0928 |  | *fepG* | Poultry and Human | *Escherichia coli*  CFT073 | Ferric enterobactin transport system permease protein fepG | Iron uptake |
| VFG0930 |  | *entF* | Human | *Escherichia coli*  CFT073 | Enterobactin synthetase component F | Iron uptake |
| VFG0931 |  | *entC* | Poultry and Human | *Escherichia coli*  CFT073 | Isochorismate synthase entC | Iron uptake |
| VFG0932 |  | *entE* | Poultry and Human | *Escherichia coli*  CFT073 | Enterobactin synthetase component E | Iron uptake |
| VFG0933 |  | *entB* | Poultry and Human | *Escherichia coli*  CFT073 | Isochorismatase | Iron uptake |
| VFG0934 |  | *entA* | Poultry and Human | *Escherichia coli*  CFT073 | "2,3-dihydro-2,3-dihydroxybenzoate dehydrogenase" | Iron uptake |
| VFG0935 |  | *iroN* | Poultry | *Escherichia coli*  CFT073 | Siderophore receptor IroN | Iron uptake |
| VFG0938 |  | *iucC* | Human | *Escherichia coli*  CFT073 | IucC protein | Iron uptake |
| VFG0939 |  | *iucB* | Human | *Escherichia coli*  CFT073 | IucB protein | Iron uptake |
| VFG0369 |  | *int* | Poultry and Human | *Yersinia pestis* CO92 | integrase | Integrase |
| VFG1028 |  | *intI1* | Poultry and Human | *Shigella* | Tn21 integrase IntI1 | Integrase |
| VFG1693 |  | *int* | Poultry and Human | *Escherichia* | Prophage P4 integrase | Integrase |
| VFG0647 |  | SF2983 | Human | *Shigella flexneri* (serotype 2a) | Transposase of Tn10 | Transposase |
| VFG0785 |  | Z5089 | Poultry | *Escherichia* | Putative transposase | Transposase |
| VFG1494 |  | s0055 | Human | *Escherichia* | Putative IS110 transposase | Transposase |
| VFG1513 |  | s0025 | Human | *Escherichia* | IS66-like transposase | Transposase |
| VFG1631 |  | *insF* | Human | *Escherichia* | Transposase of insertion element IS3 | Transposase |
| VFG1655 |  | *insG* | Human | *Escherichia* | Transposase InsG of insertion element IS4 | Transposase |
| VFG1706 |  | c3575 | Human | *Escherichia* | Transposase insF for insertion sequence IS3A/B/C/D/E/fA | Transposase |
| VFG1718 |  | c3597 | Human | *Escherichia* | Transposase | Transposase |
| VFG0477 |  | *rpoS* | Poultry and Human | *Salmonella enterica* (serovar typhimurium) LT2 | Sigma S (sigma 38) factor of RNA polymerase | Regulation |
| VFG2340 |  | *fliR* | Human | *Yersinia enterocolitica* 8081 | Flagellar biosynthetic protein FliR | Secretion system |
| VFG2055 |  | *gspL* | Poultry and Human | *Shigella dysenteriae* Sd197 | Putative general secretion pathway for protein export | Secretion system |
| VFG2046 |  | *gspC* | Poultry and Human | *Shigella dysenteriae* Sd197 | Putative general secretion protein GspC | Secretion system |
| VFG2056 |  | *gspM* | Poultry and Human | *Shigella dysenteriae* Sd197 | Putative secretion pathway protein | Secretion system |
| VFG2047 |  | *gspD* | Poultry and Human | *Shigella dysenteriae* Sd197 | Putative type II secretion protein | Secretion system |
| VFG2048 |  | *gspE* | Poultry and Human | *Shigella dysenteriae* Sd197 | Putative type II secretion protein | Secretion system |
| VFG2049 |  | *gspF* | Poultry and Human | *Shigella dysenteriae* Sd197 | Putative type II secretion protein | Secretion system |
| VFG2050 |  | *gspG* | Human | *Shigella dysenteriae* Sd197 | Putative type II secretion protein | Secretion system |
| VFG2052 |  | *gspI* | Poultry and Human | *Shigella dysenteriae* Sd197 | Putative type II secretion protein | Secretion system |
| VFG2053 |  | *gspJ* | Poultry and Human | *Shigella dysenteriae* Sd197 | Putative type II secretion protein | Secretion system |
| VFG2054 |  | *gspK* | Poultry and Human | *Shigella dysenteriae* Sd197 | Putative type II secretion protein | Secretion system |
| VFG2319 |  | *fliA* | Poultry and Human | *Yersinia enterocolitica* 8081 | RNA polymerase sigma factor for flagellar operon | Secretion system |
| VFG1871 |  | *lspG* | Poultry | *Legionella pneumophila* Philadelphia 1 | Type II protein secretion LspG | Secretion system |

**Supplementary Table 5: Samples and its detail that was collected from Thapathali temporary settlement for this study.**

| **Sample code** | **Sample type** | **Location** |
| --- | --- | --- |
| TH990 | Human faeces | Temporary settlement near Bagmati river, Thapathali |
| TH980 | Human faeces | Temporary settlement near Bagmati river, Thapathali |
| TH950 | Human faeces | Temporary settlement near Bagmati river, Thapathali |
| TH940 | Human faeces | Temporary settlement near Bagmati river, Thapathali |
| TH920 | Human faeces | Temporary settlement near Bagmati river, Thapathali |
| TH910 | Human faeces | Temporary settlement near Bagmati river, Thapathali |
| TH1150 | Human faeces | Temporary settlement near Bagmati river, Thapathali |
| TH1140 | Human faeces | Temporary settlement near Bagmati river, Thapathali |
| TH1120 | Human faeces | Temporary settlement near Bagmati river, Thapathali |
| TH1110 | Human faeces | Temporary settlement near Bagmati river, Thapathali |
| TH1040 | Human faeces | Temporary settlement near Bagmati river, Thapathali |
| TH1030 | Human faeces | Temporary settlement near Bagmati river, Thapathali |
| TH1020 | Human faeces | Temporary settlement near Bagmati river, Thapathali |
| TH1010 | Human faeces | Temporary settlement near Bagmati river, Thapathali |
| TA340 | Common quail *(Coturnix coturnix)* excrement | Temporary settlement near Bagmati river, Thapathali |
| TA330 | Common quail *(Coturnix coturnix)* excrement | Temporary settlement near Bagmati river, Thapathali |
| TA320 | Chicken (*Gallus gallus domesticus)* excrement | Temporary settlement near Bagmati river, Thapathali |
| EW070 | Water | Bagmati river, near Thapathali |
| ESD060 | River bed sediments | Bagmati river, near Thapathali |
| ES060 | Soil | Agricultural land near Thapathali temporary settlement |
| ES050 | Soil | Agricultural land near Thapathali temporary settlement |
